# Human mobility at Tell Atchana (Alalakh) during the 2nd millennium BC: integration of isotopic and genomic evidence

**DOI:** 10.1101/2020.10.23.351882

**Authors:** Tara Ingman, Stefanie Eisenmann, Eirini Skourtanioti, Murat Akar, Jana Ilgner, Guido Alberto Gnecchi Ruscone, Petrus le Roux, Rula Shafiq, Gunnar U. Neumann, Marcel Keller, Cäcilia Freund, Sara Marzo, Mary Lucas, Johannes Krause, Patrick Roberts, K. Aslıhan Yener, Philipp W. Stockhammer

## Abstract

The Middle and Late Bronze Age Near East, a period roughly spanning the second millennium BC (ca. 2000-1200 BC), is frequently referred to as the first ‘international age’, characterized by intense and far-reaching contacts between different entities from the eastern Mediterranean to the Near East and beyond. In a large-scale tandem study of stable isotopes and ancient DNA of individuals excavated at Tell Atchana (Alalakh), situated in the northern Levant, we explore the role of mobility at the capital of a regional kingdom. We generated strontium isotope data for 53 individuals, oxygen isotope data for 77 individuals, and added ancient DNA data from 9 new individuals to a recently published dataset of 28 individuals. A dataset like this, from a single site in the Near East, is thus far unparalleled in terms of both its breadth and depth, providing the opportunity to simultaneously obtain an in-depth view of individual mobility and also broader demographic insights into the resident population. The DNA data reveals a very homogeneous gene pool, with only one outlier. This picture of an overwhelmingly local ancestry is consistent with the evidence of local upbringing in most of the individuals indicated by the isotopic data, where only five were found to be ‘non-local’. High levels of contact, trade, and exchange of ideas and goods in the Middle and Late Bronze Ages, therefore, seem not to have translated into high levels of individual mobility detectable at Tell Atchana.

## Introduction

The identification of human mobility, both of groups and of individuals, has been, and remains, a topic of much discussion within archaeology. The Near East during the second millennium BC is a particularly promising arena to explore many of the questions targeting mobility patterns and effects, as it has often been discussed as an era of high levels of international connectivity in areas such as trade, diplomacy, and artistic expression, documented by both the material and textual records [1–8]. The wide-ranging social, cultural, and economic contacts of this period have long been understood to involve high levels of individual mobility on a broad scale and across a wide area, as the exchange and movement of traders, artisans, and representatives of kings is well-documented [9–13]. However, there have been limited direct studies of life history and broader demographic trends during this time period, particularly in the Levant (where much of the isotopic work done on humans has been in later periods [14–20]), limiting the degree to which this can be effectively tested, although isotopic work done in second millennium BC contexts in Egypt [21, 22], Crete [23, 24], Greece [25, 26], Anatolia [27, 28], and Arabia [29] have indicated differing levels of individual mobility ranging from populations composed primarily of local individuals to those with very high levels of non-locals.

Tell Atchana (Alalakh), located in the Amuq Valley in modern day Turkey (Fig 1) is one among many urban sites in the Middle and Late Bronze Age (MBA and LBA, respectively; ca. 2000-1200 BC) Levant that functioned as the capital of a local kingdom, characterized by complex diplomatic and international relations and frequently shifting loyalties to bigger entities of the ancient Near East [30–33]. It is therefore a prime candidate for mobility studies, as there is a high likelihood that many different individuals from a wide range of origins both passed through and settled in the city.

**Fig 1.**
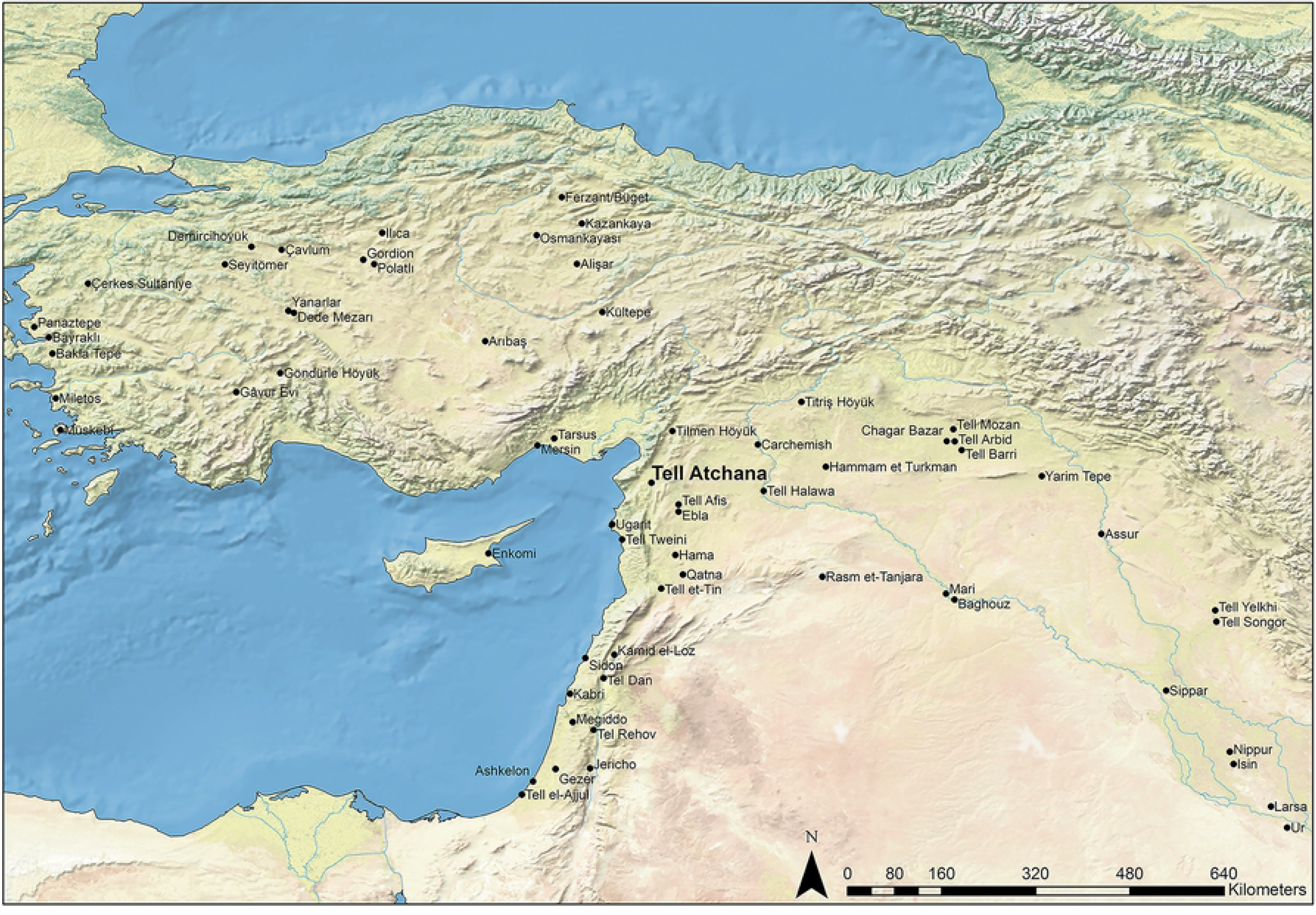
Regional map showing the location of Tell Atchana.

Isotope and ancient DNA (aDNA) analyses are two tools that shed light on the movement of individuals from different angles. With strontium and oxygen isotope ratios from tooth enamel, it is possible to identify people of non-local origin via comparison of measured ratios in the tissue of an individual and the local baseline [34–36]. Analysis of aDNA, on the other hand, sheds light on a person’s ancestry [37–39]: compared against a set of available ancient genomes of contemporary and older age from the same region and beyond, the genome of an individual holds key information about locality in terms of genetic continuity or discontinuity in an area through time or in terms of mobility as represented by genetic outlier individuals. While stable isotope analysis has been utilized in archaeology since the 1970s [34, 40, 41], full genome aDNA analyses on a large scale only became available during the last decade [37, 42]. Independently, both methods have proven powerful tools in detecting human mobility and to operate independently from archaeological concepts of burial traditions, but the exploration of their tandem potential has only recently started [43–45]. Nevertheless, the combination of both methods has yet to be applied systematically in the Ancient Near East.

In this study, we seek to explore human mobility at Tell Atchana on the basis of the most direct source available, the human remains themselves. In order to explore patterns of mobility among the individuals recovered, we performed strontium and oxygen isotope analysis and aDNA analysis on bones and teeth of individuals excavated at Tell Atchana from 2003-2017. We publish here the first strontium and oxygen isotope data of 53 and 77 individuals, respectively, and add genome-wide data for nine individuals to an existing dataset of 28 individuals recently published by Skourtanioti et al. [46], with sampled individuals coming from a wide range of different contexts. With this extensive, in-depth analysis of a large number of individuals from a single site, a study thus far unique for the ancient Near East, we demonstrate how isotope and aDNA data can complement or even contradict each other, and how both strands of evidence can be combined with the archaeological context in order to address questions regarding the nature and scale of individual mobility in the Near Eastern Bronze Age.

### Tell Atchana

Situated on the southward bend of the Orontes River in the modern state of Hatay, Turkey (see Fig 1), Tell Atchana (Alalakh) was founded in the terminal Early Bronze Age or the earliest MBA (ca. 2200-2000 BC), flourishing throughout the MBA and LBA until its nearly complete abandonment ca. 1300 BC [31–33, 47]. The site was first excavated in the 1930s-1940s by Sir Leonard Woolley [30, 48], who exposed large horizontal swathes of what came to be known as the ‘Royal Precinct’ of the site (Fig 2) and uncovered a continuous sequence of 18 levels from Level XVII to Level O [30], the latter now known to date to the Iron Age (Table 1) [32, 47, 49]. K. Aslıhan Yener returned to the Amuq Valley in 1995 [50] and resumed ongoing excavations at Tell Atchana in 2003 [31, 32].

**Fig 2.**
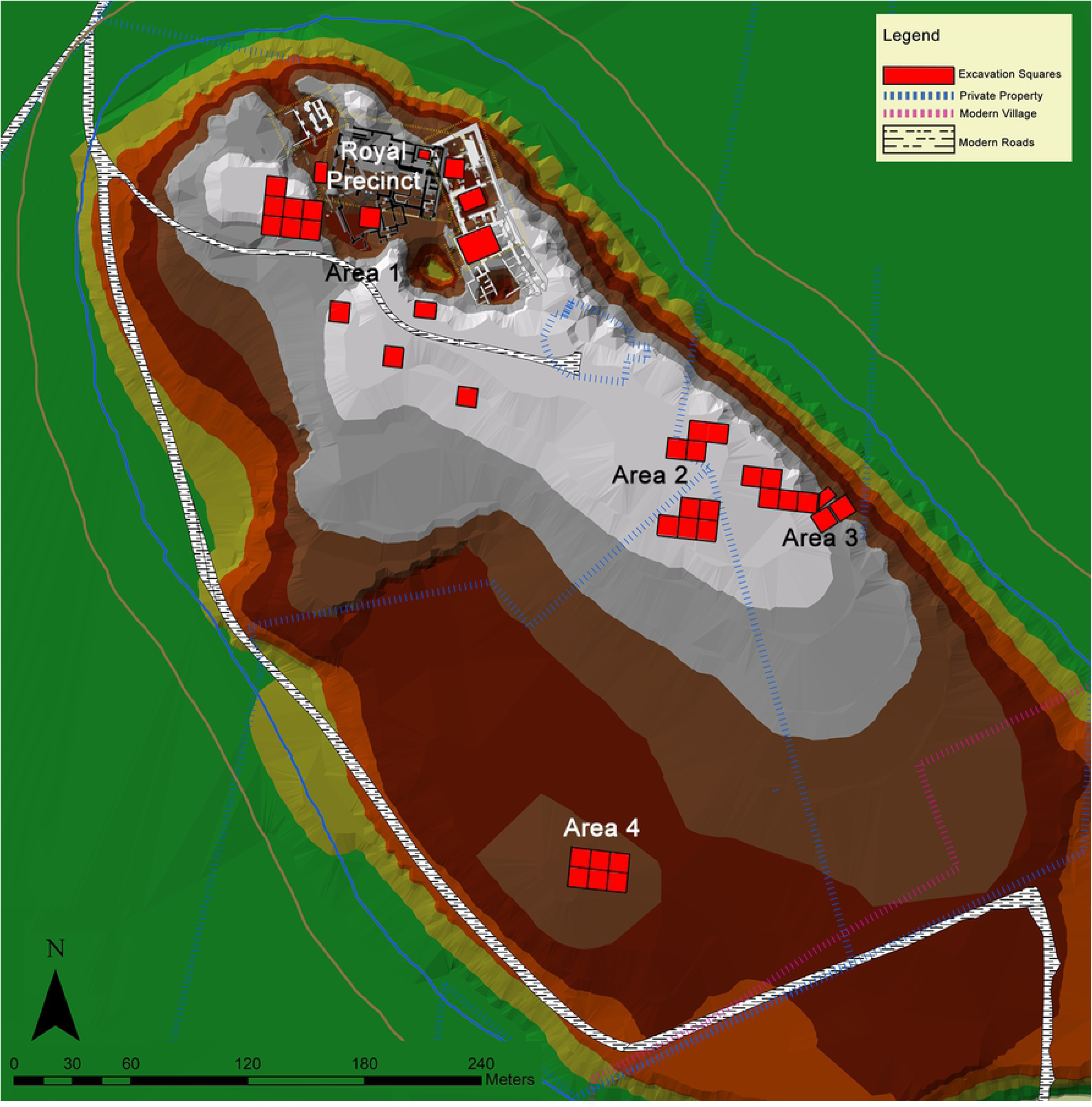
Map of Tell Atchana with excavation squares indicated.

**Table 1.**
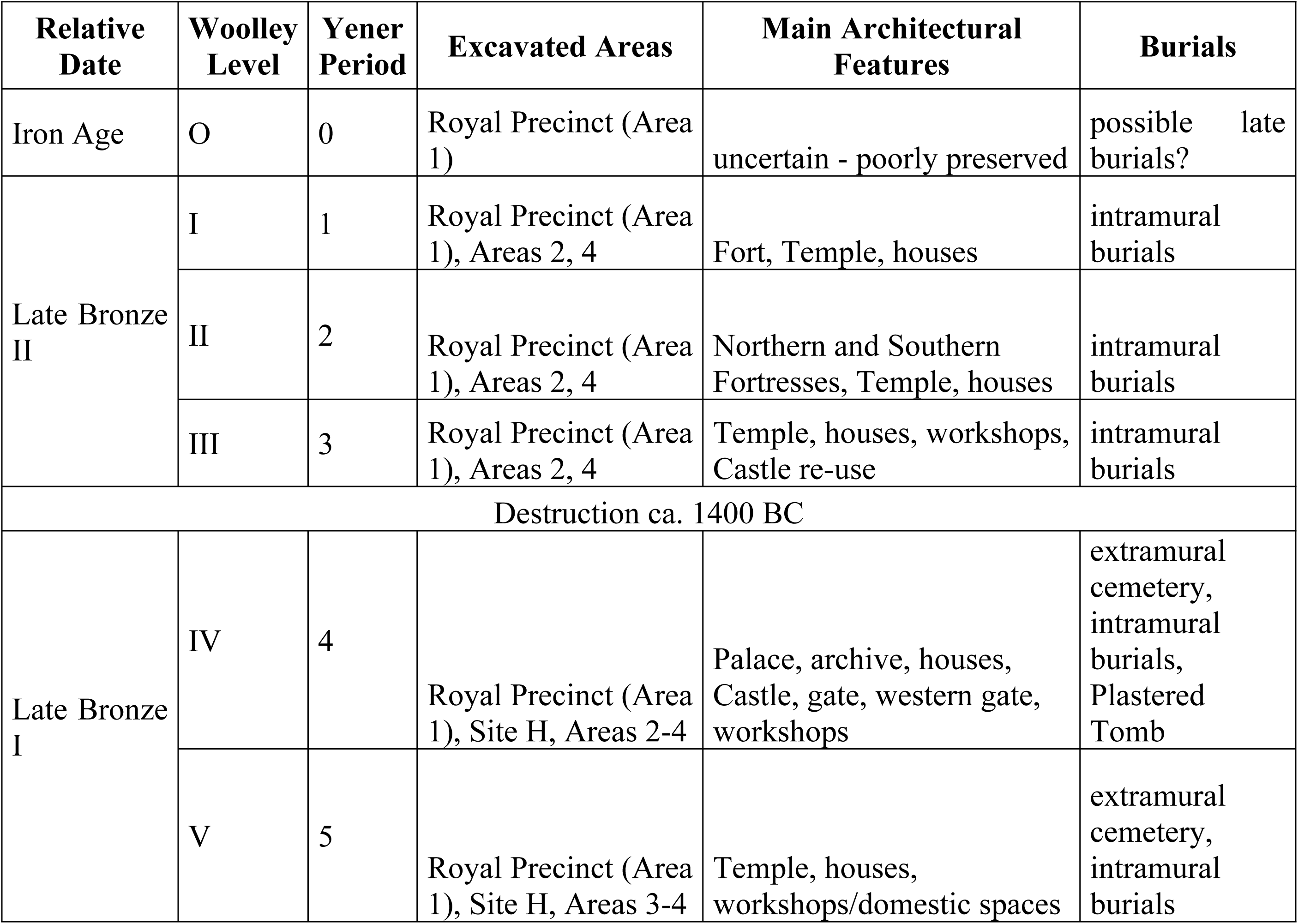
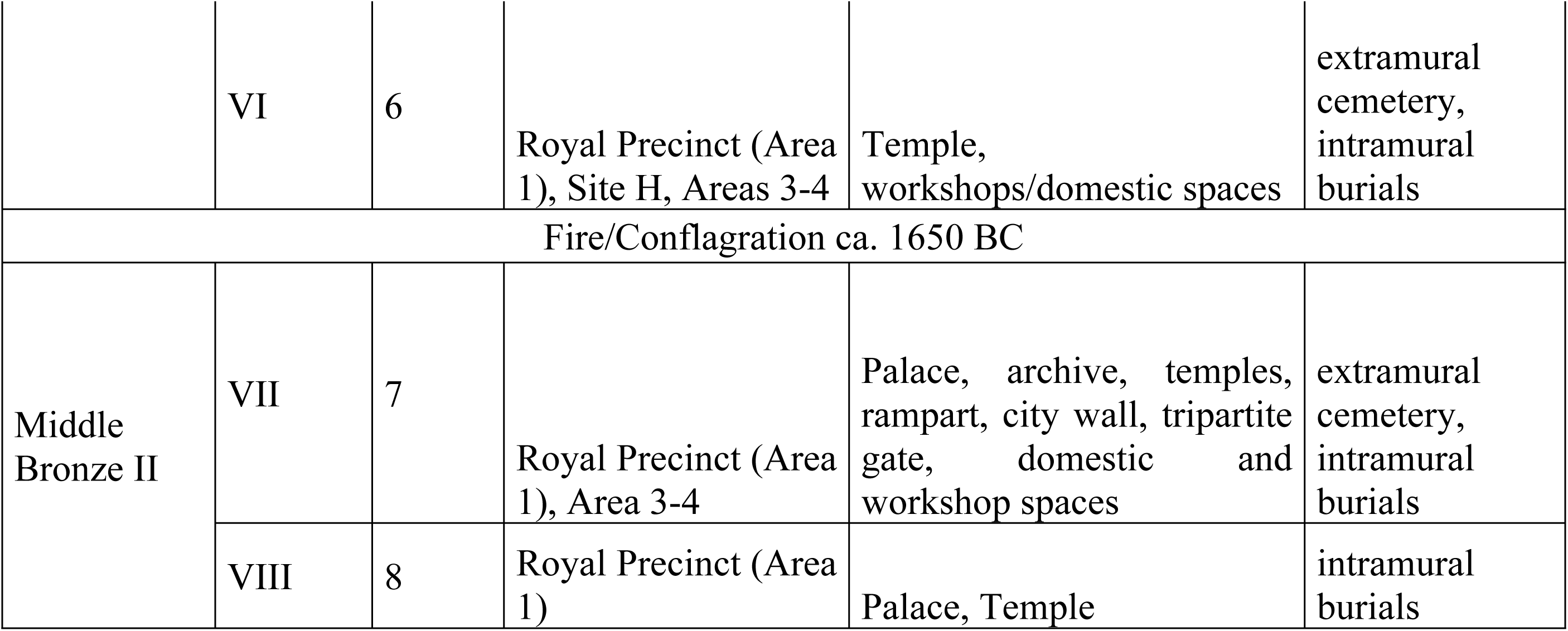
Chronology of Tell Atchana.

Texts from the palace archives dating from the MB II and LB I at Tell Atchana itself and from other sites that mention the city of Alalakh provide ample evidence about the city’s significance as the capital of the region and its relations of exchange with its neighbors, such as Ebla, Ugarit, Halab, Emar, and cities in Cilicia, as well as entities located further away, like the state of Mitanni, Mari, the Kassite kingdom of Babylonia, the Hittites, and Middle and New Kingdom Egypt [5, 31, 51–57]. The textual record is matched by an archaeological record, particularly for the LBA, rich in imports (or objects imitating foreign styles) and architecture bearing foreign influences, including particular building methods, imported ceramic styles and small finds, and artistic motifs, such as Aegean-style bull-leaping scenes [30-33, 47, 57-75]. It is unclear how strongly this evidence was connected with the actual presence of people from abroad in permanent residence at Alalakh, however. While it is likely that at least some migrants lived and died at the site, it is impossible to make claims about the actual scale on the basis of texts and archaeology alone. It is also unclear whether these migrants were buried in the 342 graves which have been excavated to date, making the site a perfect candidate for targeted mobility studies.

## Materials and methods

### Tell Atchana burial corpus

Burials at the site are present from the late MBA through the end of the LBA (stratigraphically, in contexts from Periods 8-1; see Table 1) and have been found in every excavated area of the site. Tell Atchana has one of the largest numbers of recorded burials in the area, incorporating different types of burials, burial goods, and burial locations, including both intramural burials (208 examples in total) and an extramural cemetery outside the city fortification wall in Area 3 (134 burials; see Fig 2) [76, 77]. The term ‘intramural’ is used here to differentiate these burials from the extramural burials and indicate their location within the walls of the city, rather than their location within buildings *per se*: they have been found in various contexts, such as in courtyards and other open spaces, in the ruins of abandoned buildings, and under intact floors. A total of 28 have been found in what appears to be an intramural cemetery recently discovered in the south of the mound in Area 4 (see Fig 2) [77, 78]. The presence of both intramural and extramural burials provides a rare opportunity to compare the two funerary practices at a single site.

The vast majority of the burials are single, primary pit graves, although there are a handful of secondary and/or multiple burials, as well as cist graves, pot burials, and cremations [77, 79]. This variety is a starting place to look for the presence of non-locals, who could be associated with these minority types of burials. In the extramural cemetery, grave goods are rare, with over half of the burials containing no grave goods, but when they are present, they typically consist of one or two vessels and perhaps an article of jewelry, most often a metal pin or a beaded bracelet/necklace [76]. The intramural burials, particularly those found in the Royal Precinct, are generally the richest in grave goods, with a wide variety of imported and local pottery, metal jewelry, and rarer items such as figurines and stone vessels [77, 79], supporting the suggestion that these burials represent a higher social class than the individuals interred in the extramural cemetery [59, 76, 77, 79]. The exception to this, and the most intriguing burial at the site, is the Plastered Tomb. Located in the extramural cemetery, it was built of several layers of plaster encasing four individuals that dates to the end of LB I [80–82]. This is the richest burial found at the site, with 13 vessels and numerous items of adornment, including beads made of gold, carnelian, and vitreous materials, pins of bronze and silver, and pieces of foil and stamped appliques made of gold. Due to its unique status, its unusual construction, and its rich assemblage of objects, it was a particular target for this study.

In addition to these broad burial groupings, several individuals have been recovered who seem to have died as a result of some type of misadventure and did not receive formal burials, two of which are included in this study. The first, the so-called ‘Well Lady’ (ALA019), whose skeletal remains were found at the bottom of a well, was apparently thrown into the well while it was still in use, and homicide has been proposed as her manner of death [83]. The second, an adult female (ALA030), seems to have been killed during the destruction and collapse of a building in Area 3 [84].

#### The chronology of the burials

The ^14^C-AMS-dating published in Skourtanioti et al. [46] included 21 individuals from the extramural cemetery (Table 2, Fig 3). It indicates that the beginning of the cemetery’s use dates back into the MB I (i.e. before 1800 BC) and makes the extramural cemetery one of the oldest features that has been excavated at Tell Atchana to date. Furthermore, the radiocarbon dates of the extramural cemetery show a general discrepancy with the archaeological dating: while the former suggests that all individuals sampled (with the exception of those in the Plastered Tomb) date to the MBA (before 1600 cal BC), the latter puts the main use of the cemetery into LB I (ca. 1600-1400 BC), with only very few burials dated to MB II (ca. 1800-1600 BC) [76]. The reasons for this discrepancy could be general errors in the calibration curve for the Levantine area and/or that parts of the cemetery were only used during the MBA. It seems rather unlikely that by chance only those extramural cemetery individuals which belong to the MBA were radiocarbon dated (for a detailed discussion of the dates and the stratigraphy see S1 Text). Compared to the ^14^C-results from the extramural cemetery, the dates from the intramural burials show a higher level of concordance with the archaeological (stratigraphic) dating, with only two out of eight ^14^C dates being substantially earlier (ALA016 and ALA020).

**Fig 3.**
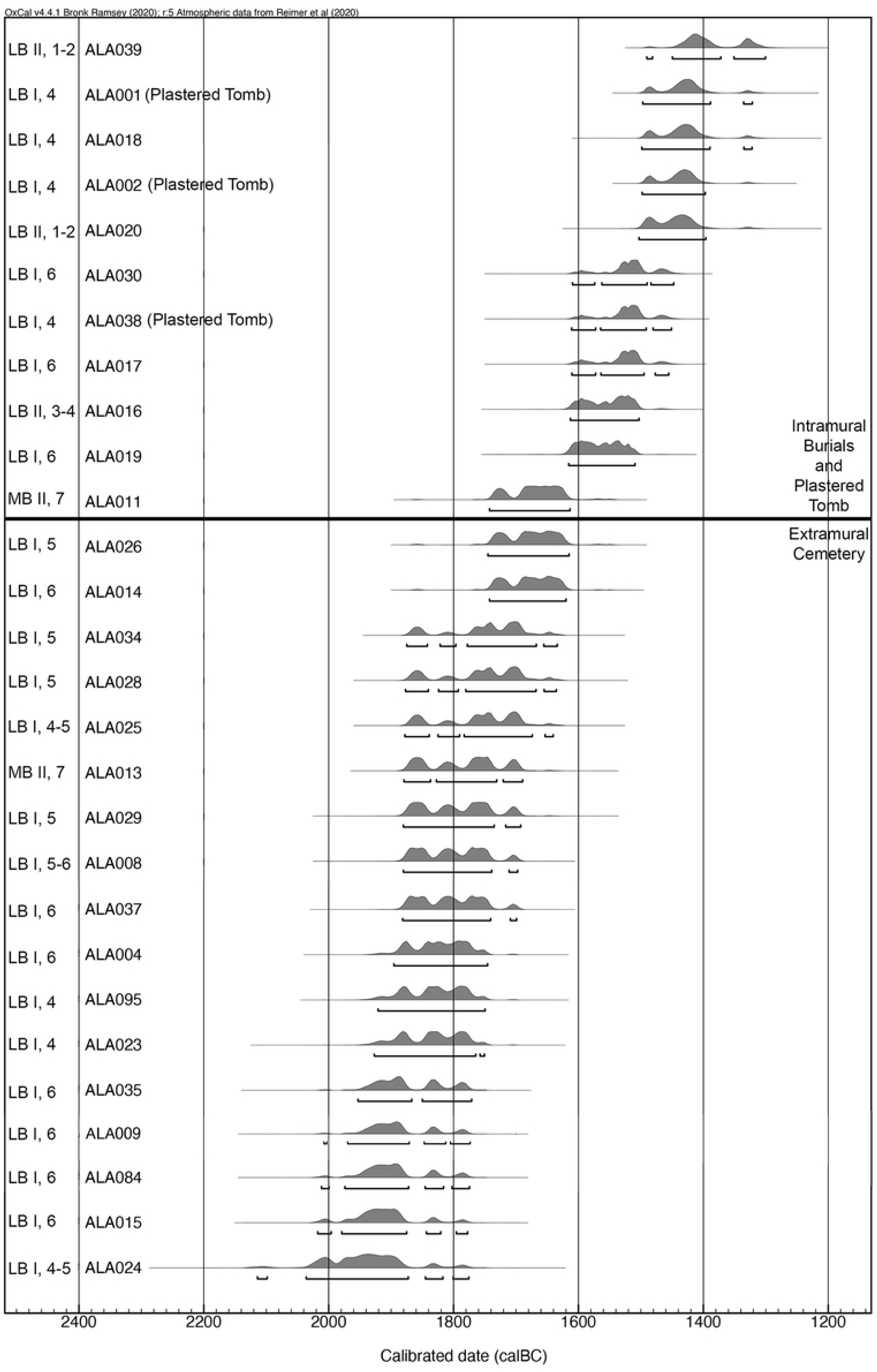
All ^14^C dates from burials at Tell Atchana, including tentative archaeological dating to Period and relative archaeological era (indicated as [ERA], [PERIOD] to the left of the individuals sample IDs).

**Table 2.**
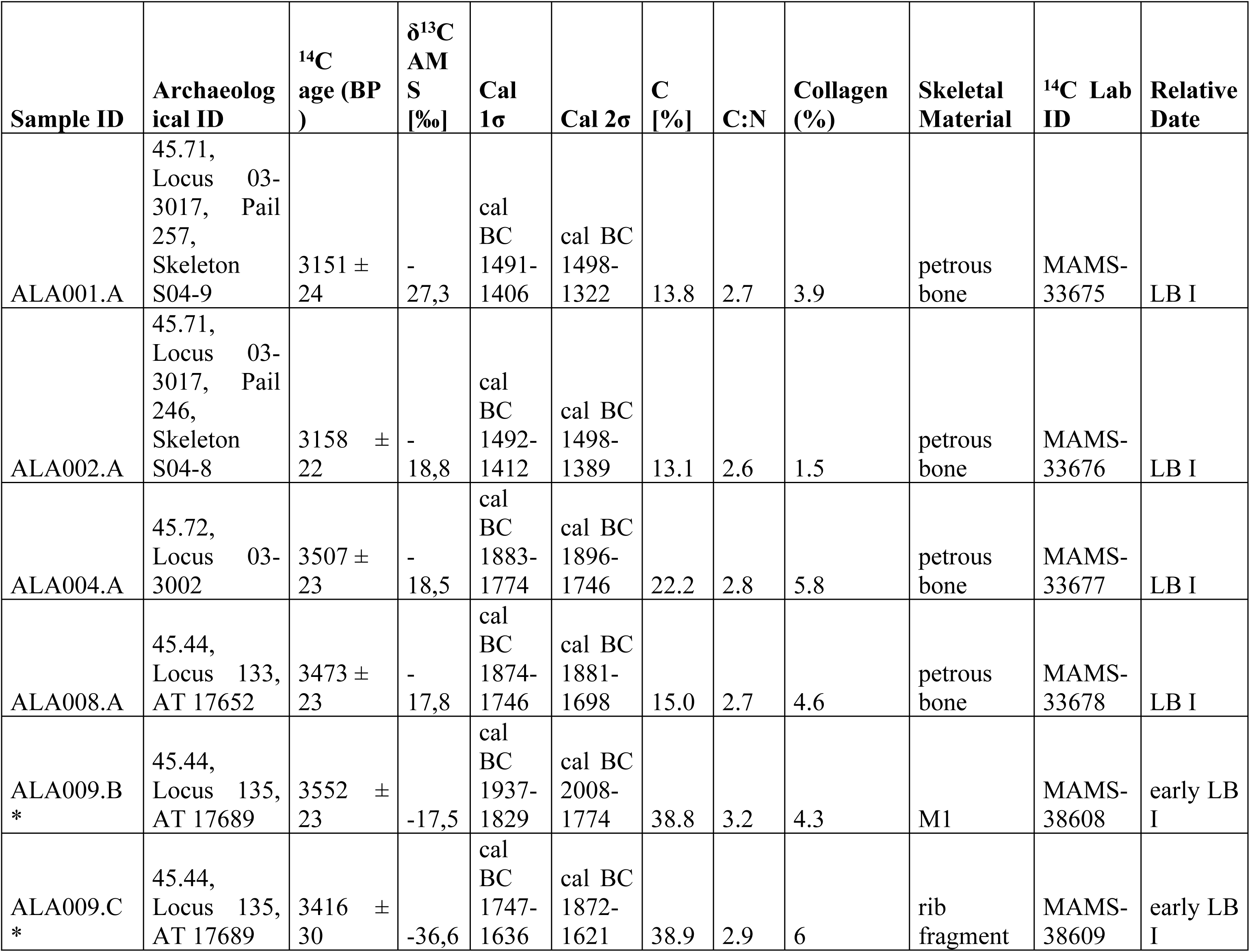
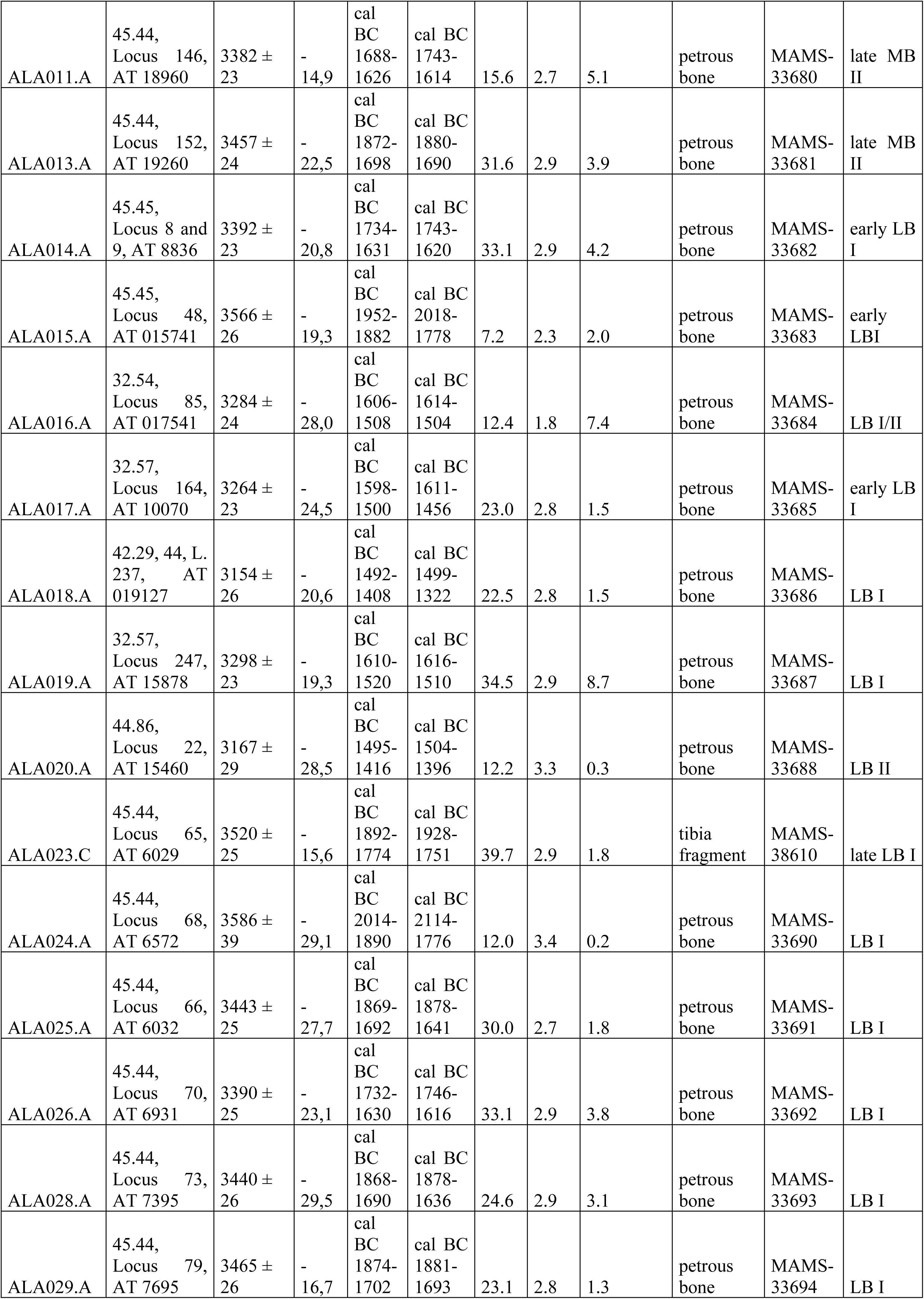
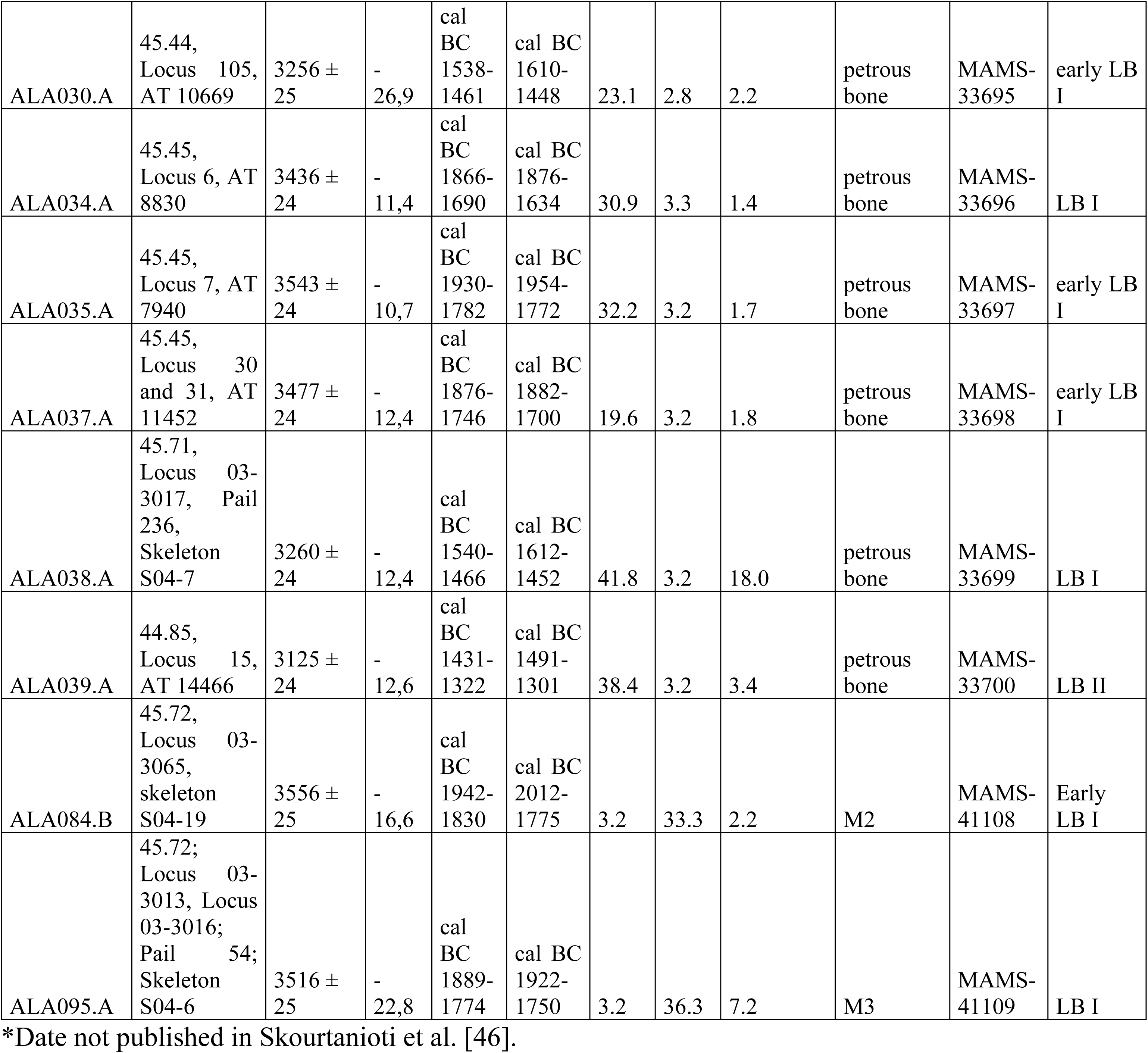
All ^14^C dates from individuals, first published in Skourtanioti et al. [46].

#### Sampling strategies and the datasets

Individuals for aDNA and isotope sampling were selected in order to be as representative as possible of the burial corpus as a whole, choosing individuals from all available intra- and extramural contexts, different types of burials (primary and secondary, single and multiple), varying age groups (with an emphasis on adult individuals), and both sexes, with age and sex data based on osteological analysis conducted by R. Shafiq. For aDNA analysis, we primarily targeted the petrous bone, the skeletal element which has been shown to best preserve human DNA, and as a secondary potential element, we used teeth [85–87]. For isotope analysis, we preferentially used permanent second molars, as the M2 is formed between the ages of ca. 2-8 years [88, 89], thereby being more likely to show isotopic signals with minimal interference from breastfeeding effects [27, 90–92]. Where no second molar was available, the M3 (formed between ca. 7-14 years [88]), M1 (formed between ca. the last month in utero to 3 years of age [88]), or a premolar (formed between ca. 1-7 years, depending on which premolar [88]) were sampled in descending order of preference. Environmental bulk reference samples (n = 16) for isotopic analysis were taken from modern and archaeological snails, as well as archaeological rodents (Table 3), in order to establish a local range for biologically available strontium, both at Tell Atchana and across the Amuq Valley more broadly. Five bulk faunal samples were also taken for oxygen isotopic analysis in order to compare the results to those of the humans.

**Table 3.** All faunal samples.

| Sample ID | $^{87}\text{Sr}/^{86}\text{Sr}$ | $\pm 2$ SD internal | $\delta^{18}\text{O}$ (‰) | SD | Species | Context |
| --- | --- | --- | --- | --- | --- | --- |
| AT 0263 | - | - | -2.2 | 0.05 | *Bos taurus* | Sq. 64.82.2 |
| AT 1074 | - | - | -7.2 | 0.04 | *Bos taurus* | Sq. 64.82.17 |
| AT 1141 | 0.708544 | 0.000010 | -5.3 | 0.16 | *Spalax leucodon* | Sq. 64.72.8 |
| AT 11570 | 0.708440 | 0.000009 | - | - | *Gastropoda* | Sq. 42.29.9 |
| AT 12146 | 0.708411 | 0.000013 | - | - | *Gastropoda* | Sq. 32.54.66 |
| AT 12952 | 0.708296 | 0.000010 | - | - | *Gastropoda* | Sq. 32.57.219 |
| AT 2051 | 0.708111 | 0.000015 | - | - | *Gastropoda* | Sq. 33.32.1 |
| AT 3061 | 0.708418 | 0.000012 | - | - | *Rodentia* | Sq. 64.73.9 |
| AT 3064 | - | - | -2.4 | 0.03 | *Caprinae* | Sq. 64.73.7 |
| AT 8302 | - | - | -4.4 | 0.05 | *Sus scrofa* | Sq. 64.82.56 |
| AT 9580 | 0.708305 | 0.000011 | - | - | *Gastropoda* | Sq. 45.44.94 |
| G1.5A | 0.708807 | 0.000013 | - | - | modern<br>*Gastropoda* | Kamberli |
| G2.2 | 0.707924 | 0.000013 | - | - | modern<br>*Gastropoda* | Kırıkhan |
| G2.6 | 0.707984 | 0.000012 | - | - | modern<br>*Gastropoda* | Reyhanlı |
| G3.3B | 0.708359 | 0.000013 | - | - | modern<br>*Gastropoda* | Hacıpaşa |
| G3.4A | 0.708661 | 0.000011 | - | - | modern<br>*Gastropoda* | Hacıpaşa |
| G4.2C | 0.708302 | 0.000011 | - | - | modern<br>*Gastropoda* | UyduKent |
| G5.5 | 0.708522 | 0.000014 | - | - | modern<br>*Gastropoda* | Avcılar |
| G6.2A | 0.708211 | 0.000015 | - | - | modern<br>*Gastropoda* | Ceylanlı |
| G6.7 | 0.707376 | 0.000014 | - | - | modern<br>*Gastropoda* | Haydarlar |

Analysis of aDNA – which, as an organic material, is subject to *post-mortem* decomposition – has a variable success rate: samples from 116 individuals from Alalakh were analyzed, but 1240K SNP data could be produced only for 37 (including both this study and Skourtanioti et al. [46]). An “ALAXXX” sample number was assigned to each analyzed individual (Table 4). All samples were photographed and documented prior to any destructive sampling, and teeth were additionally CT scanned at Max Planck Institute for the Science of Human History (MPI-SHH) in order to preserve a complete record of dental features. Currently, ^87^Sr/^86^Sr results from tooth enamel samples are available for 53 individuals, δ^18^O results for 77 individuals, and aDNA results for 37 individuals (see Table 4; see also S1 Table).

**Table 4.**
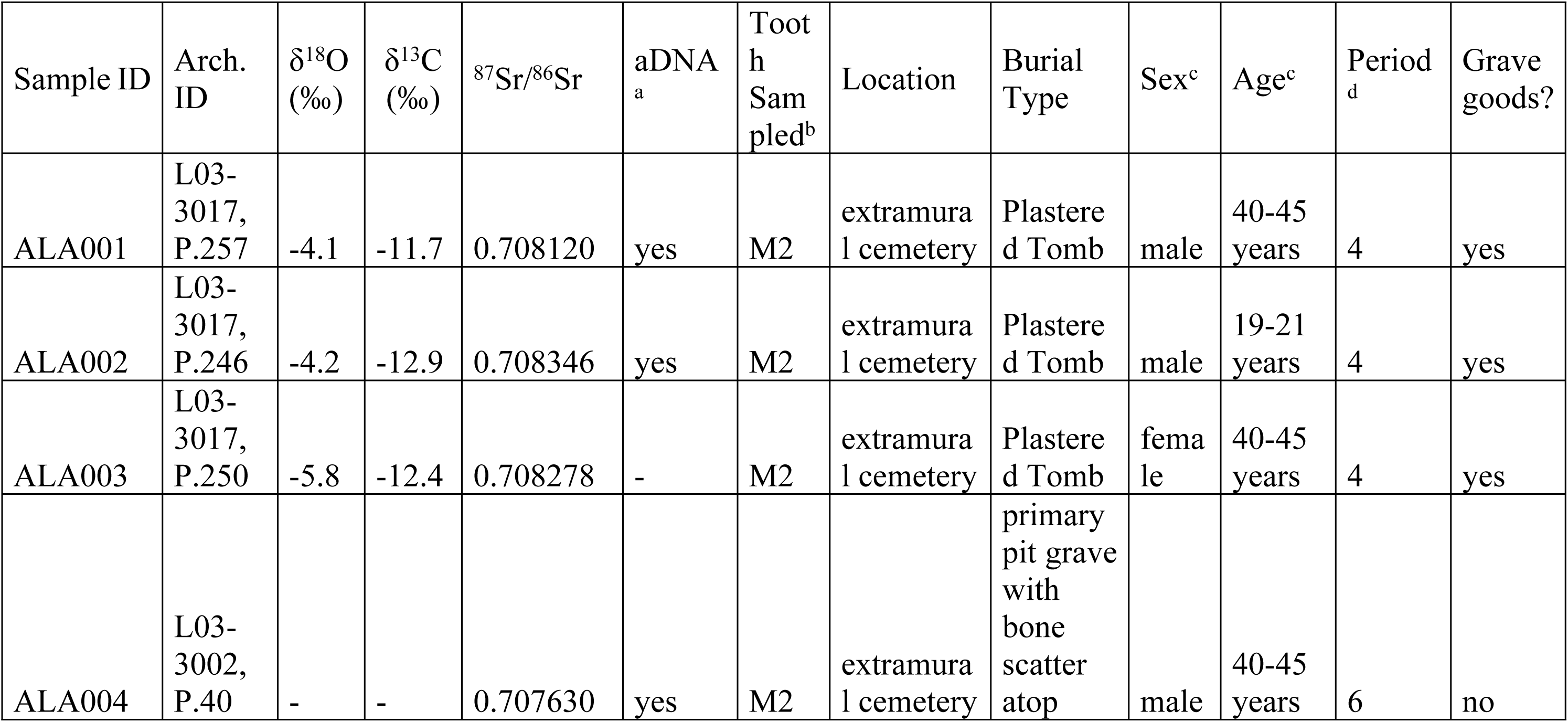
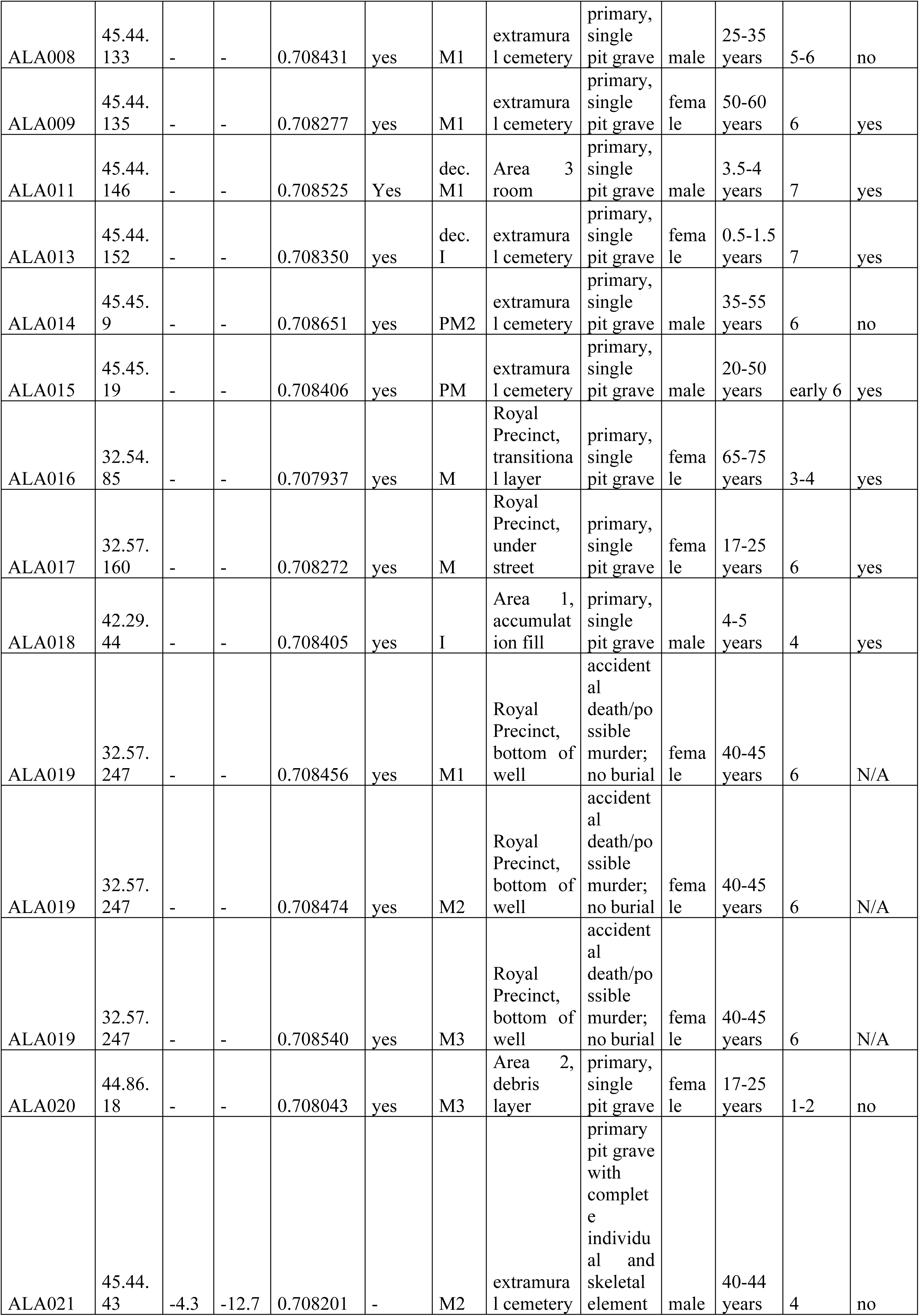
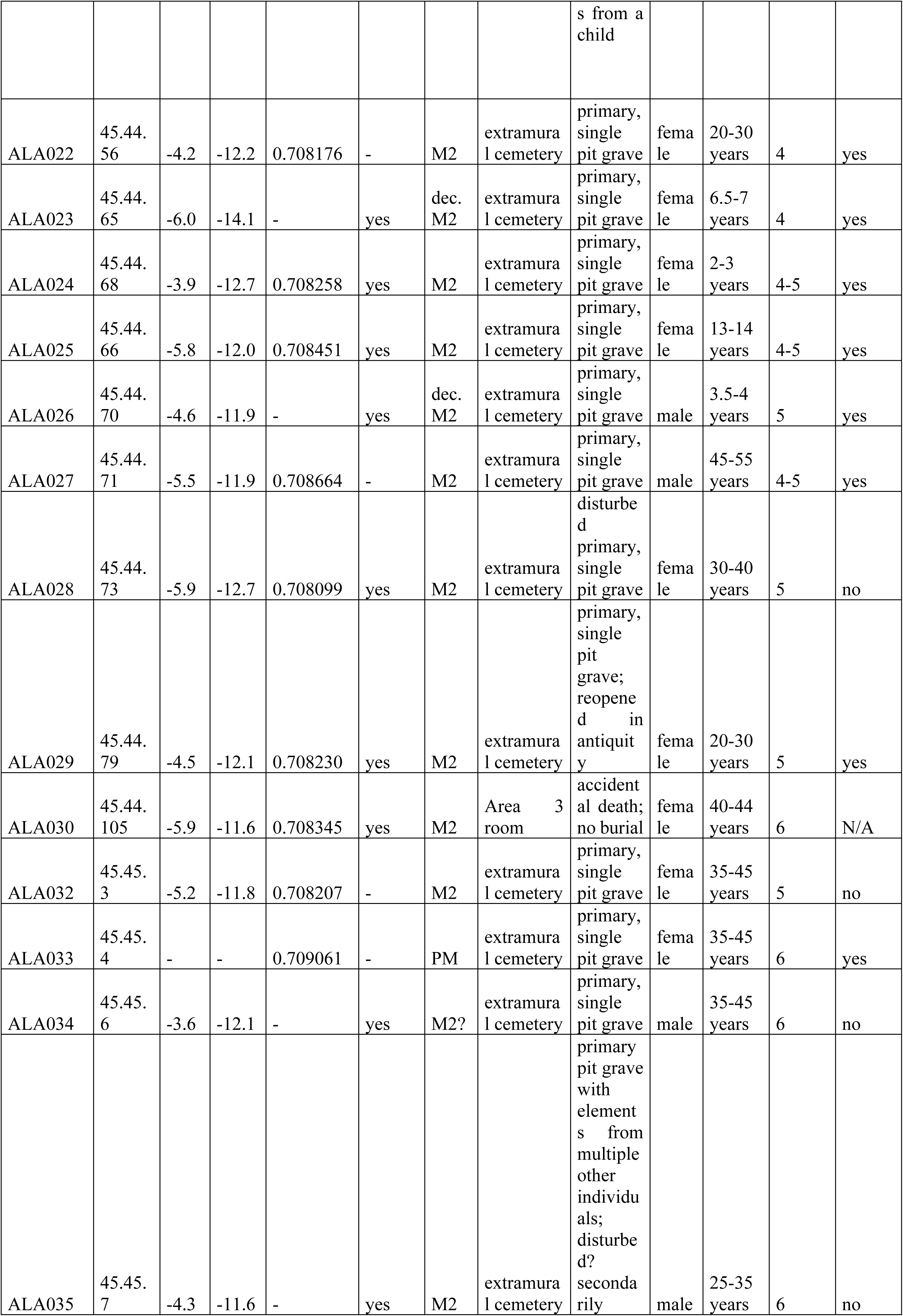
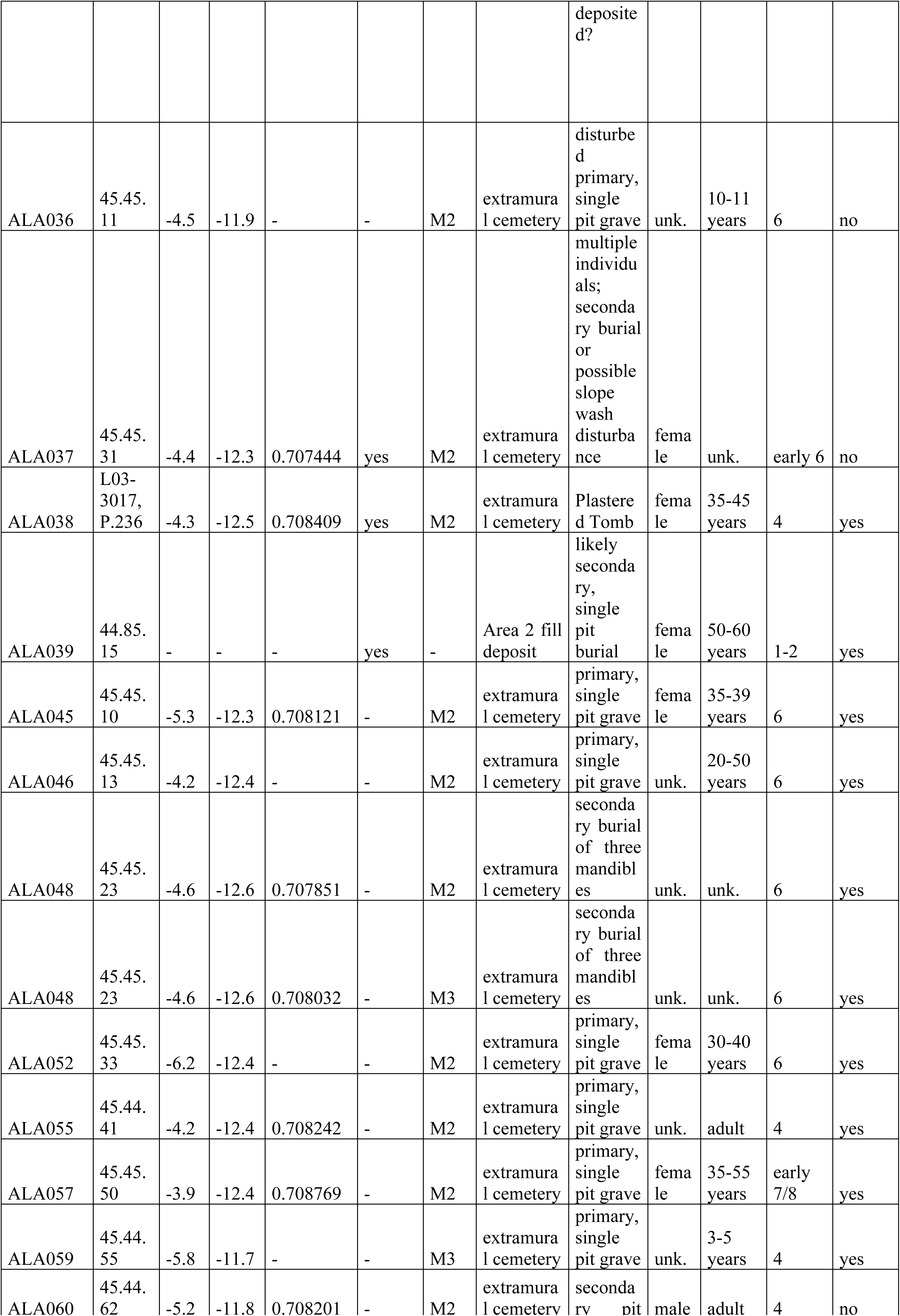
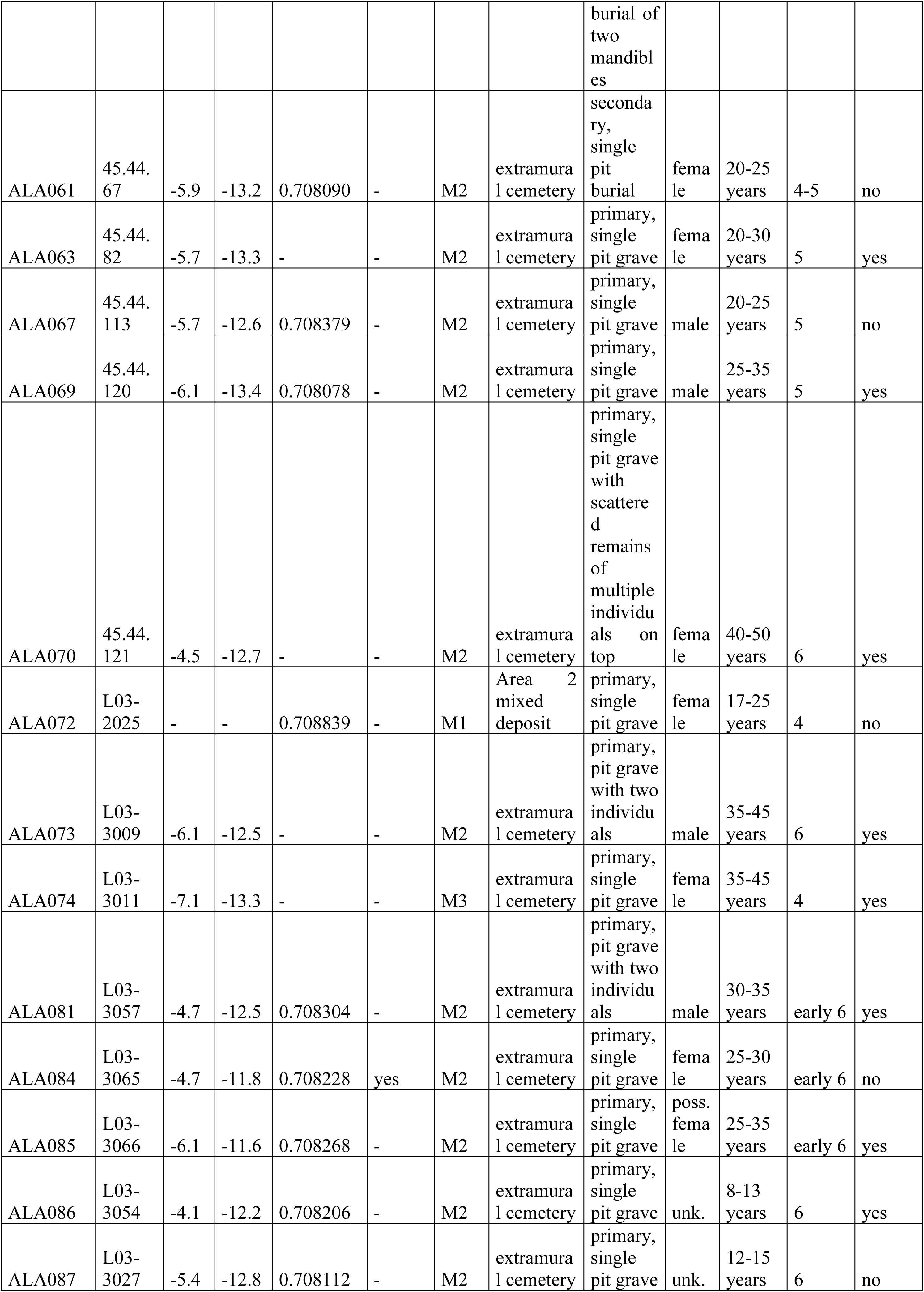
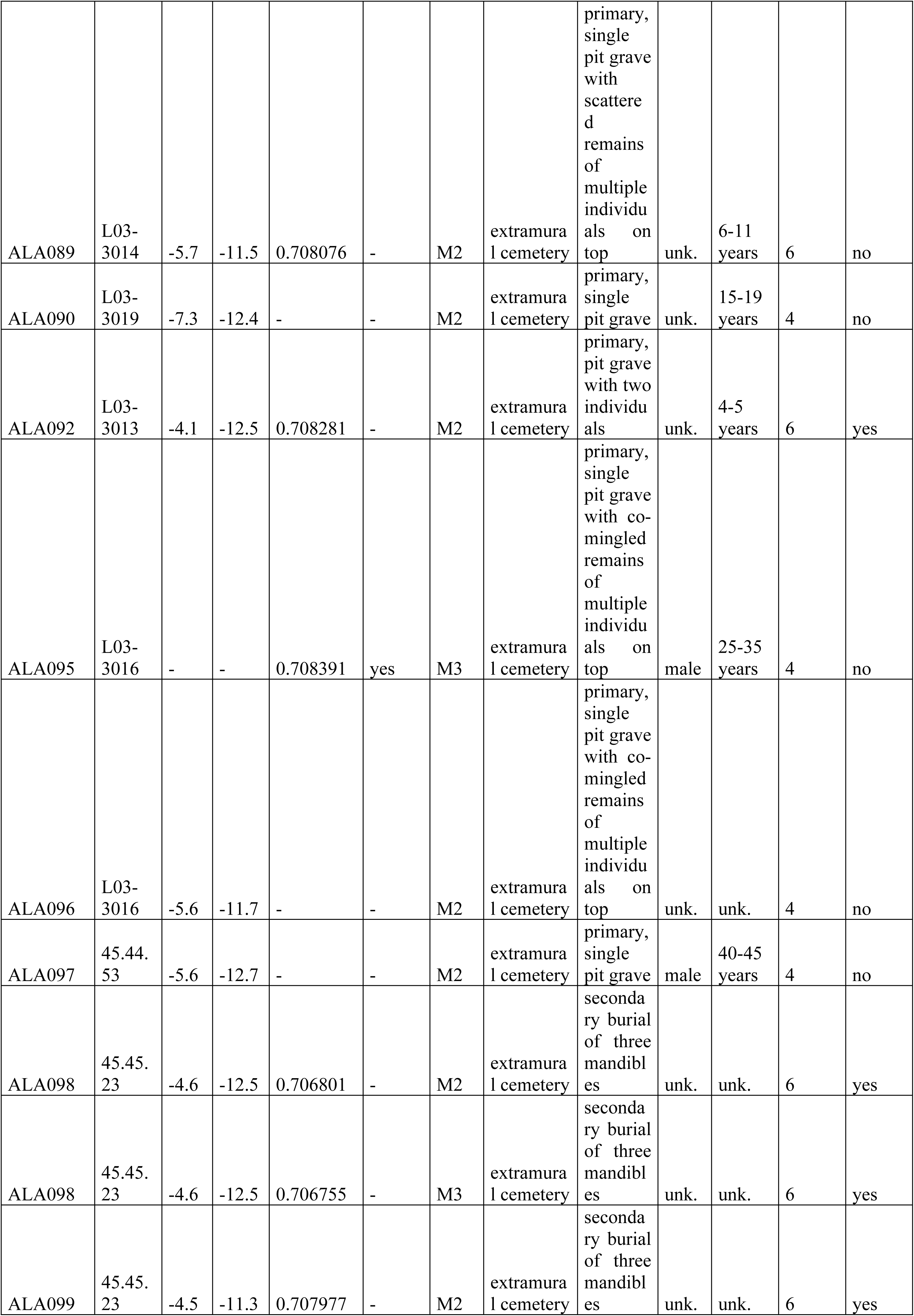
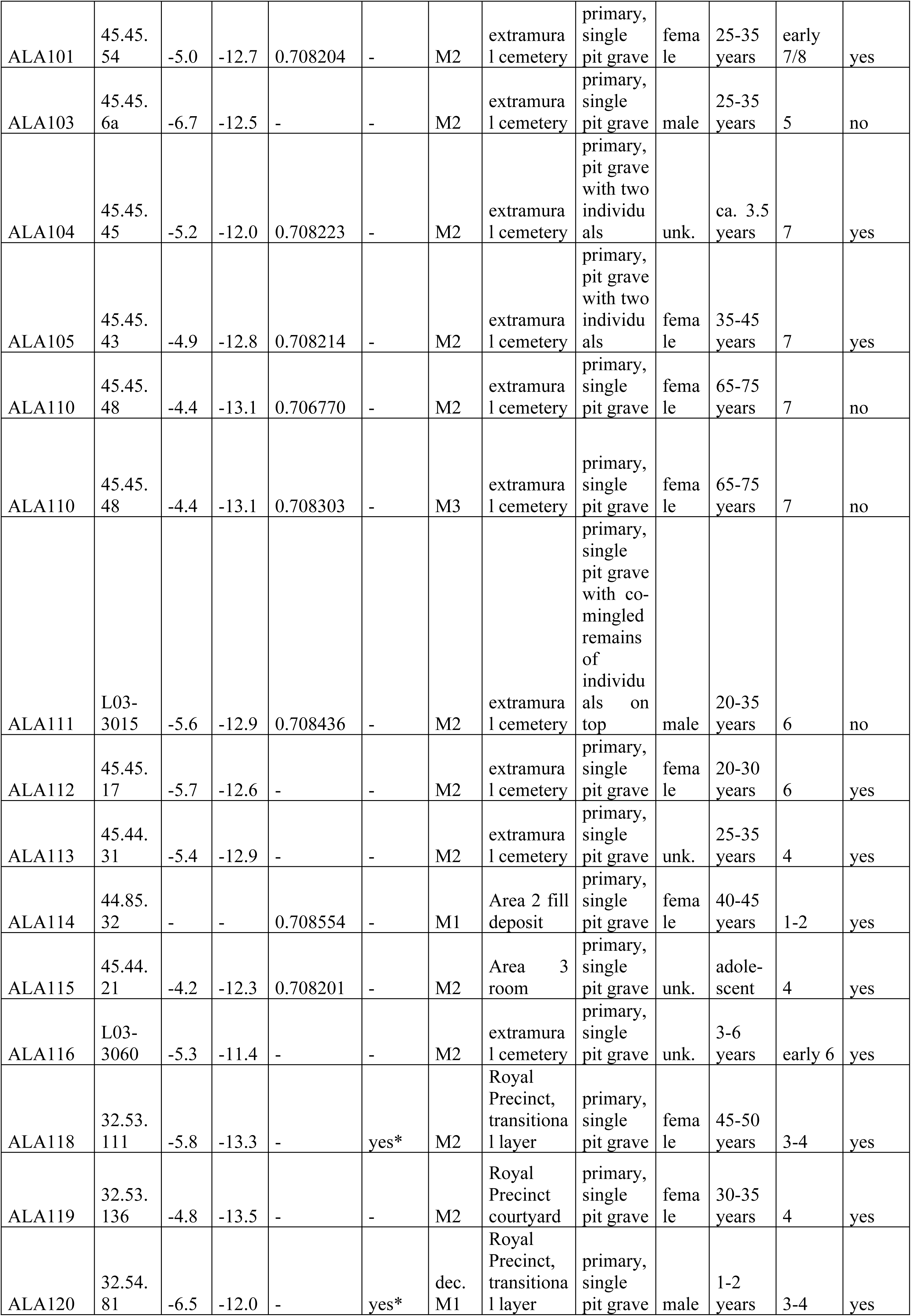
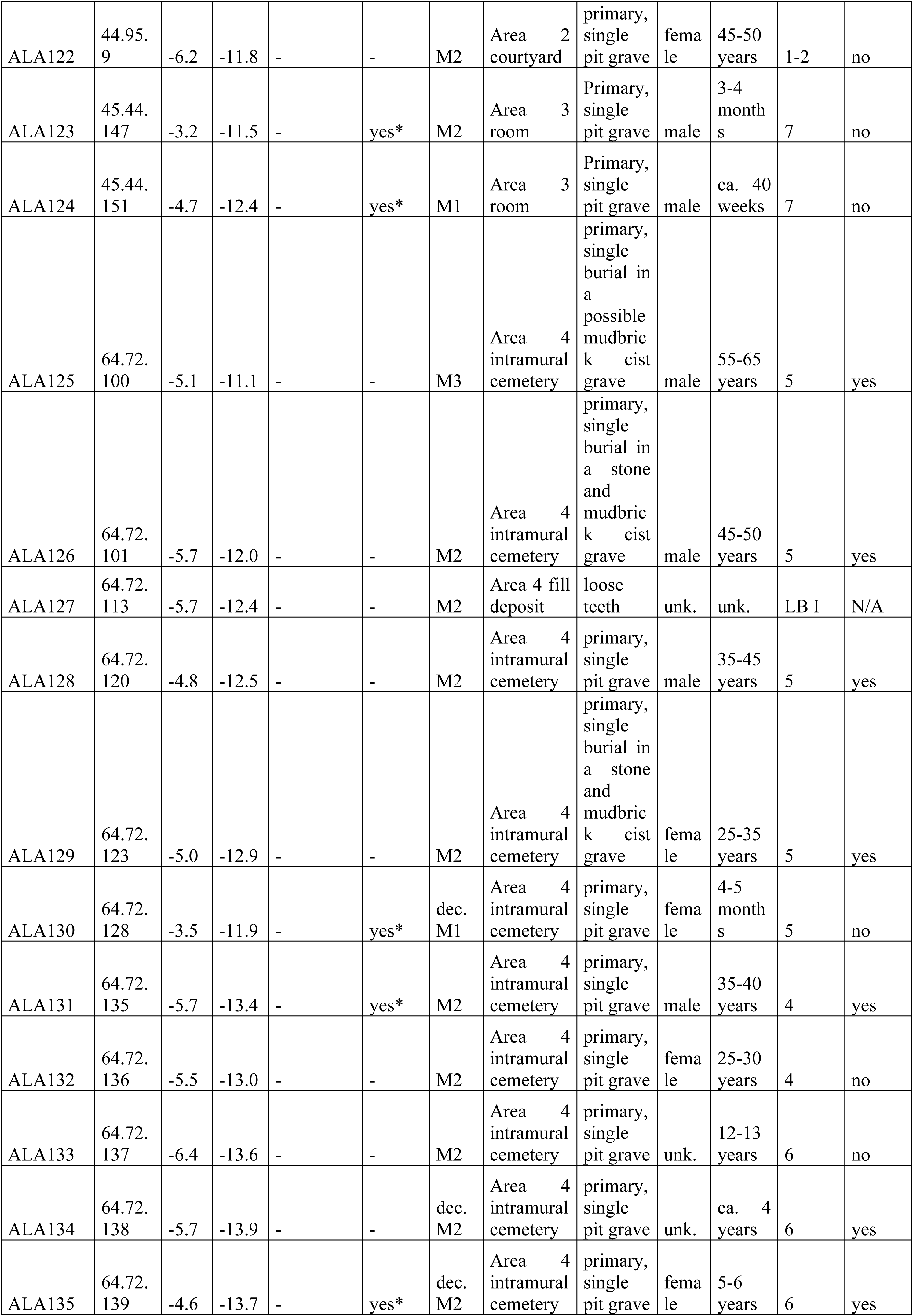
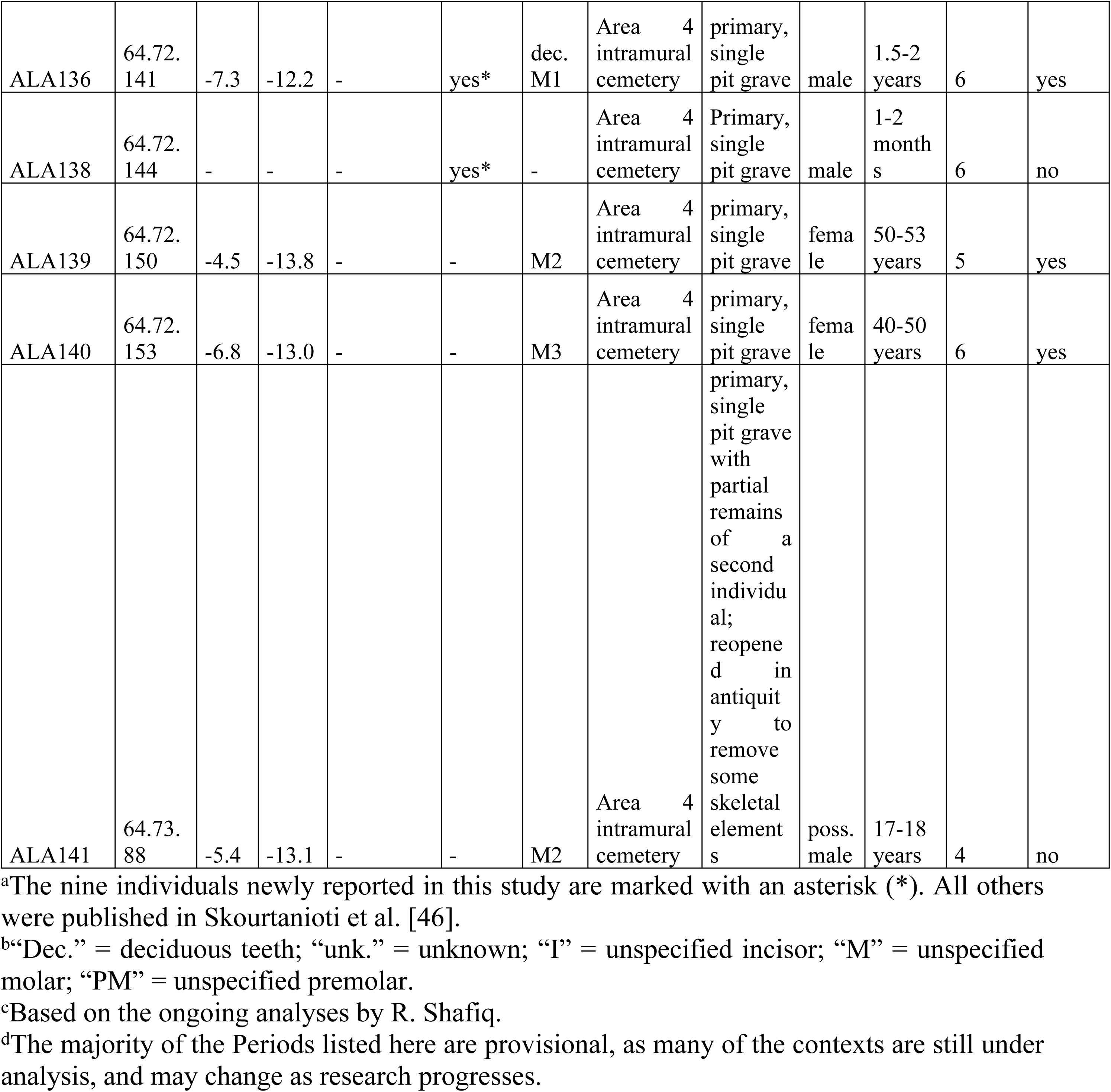
All individuals included in this study.

Although the sampled skeletal assemblage does not reflect the excavated burials at Alalakh as a whole, as the sampled individuals are biased towards the extramural cemetery (Fig 4), it includes individuals from all Areas excavated by Yener. This imbalance is a result of the fact that nearly three-quarters of the intramural burials were recovered during the previous excavations by Woolley (151 individuals of 208 intramural burials = 72.6%) and are therefore unavailable for sampling, as Woolley did not keep the human remains he found (see Fig 4). The situation is similar for the numbers of individuals sampled from each archaeological period (Fig 5), as the majority of the LB II individuals were excavated by Woolley [77]. Sub-adults as a group generally are also somewhat underrepresented among the analyzed individuals (Fig 6), due to this study’s preference for 2^nd^ (and 3^rd^) permanent molars, but the proportions of age classes is, again, roughly representative of the available material. Given the limitations of available material, therefore, the sampled individuals are as representative as possible for the excavated burials as a whole, and, most importantly, cover all known contexts and burial types.

**Fig 4.**
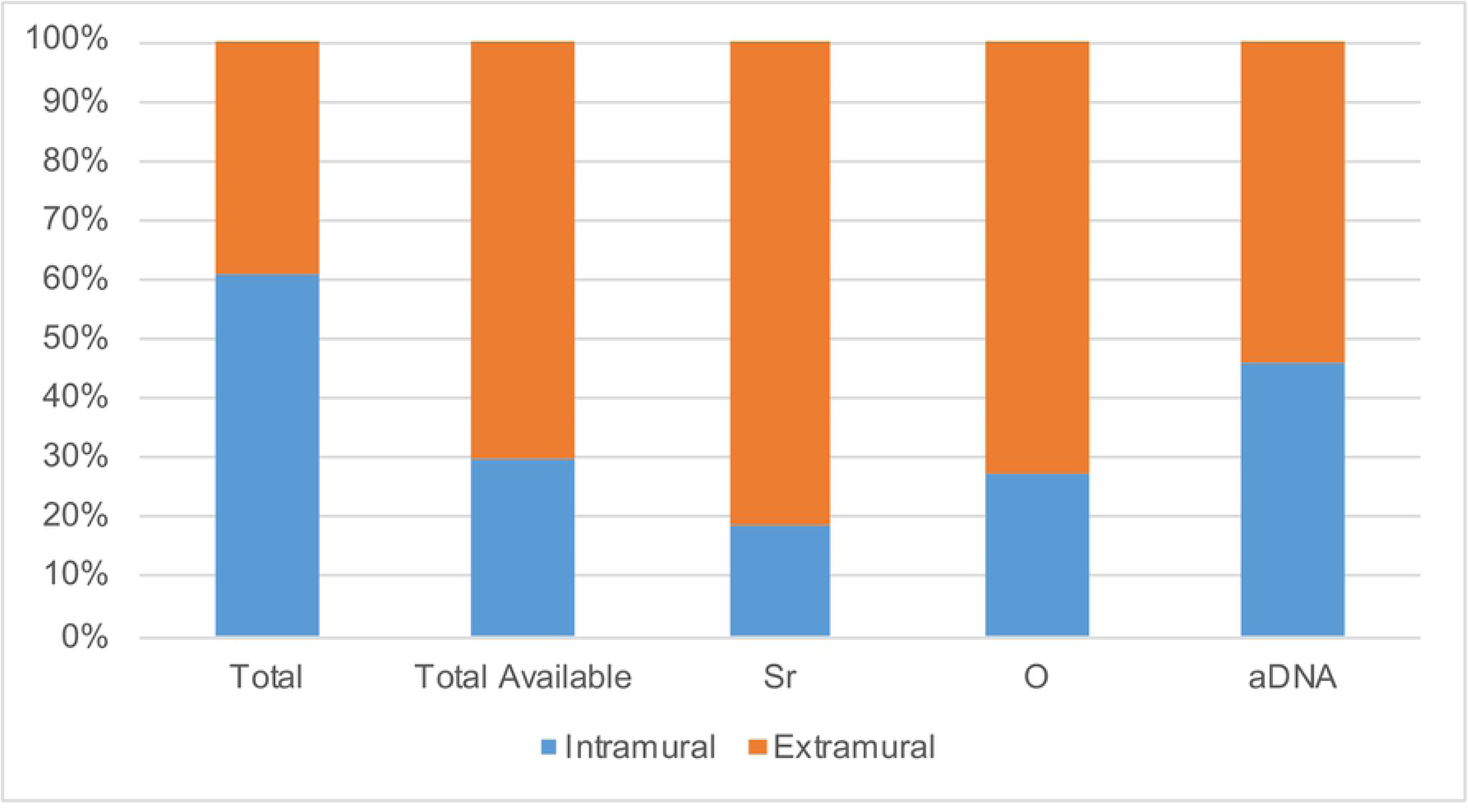
Contexts of the total assemblage available for sampling (i.e., excavated by Yener), as well as those sampled for each analysis presented.

**Fig 5.**
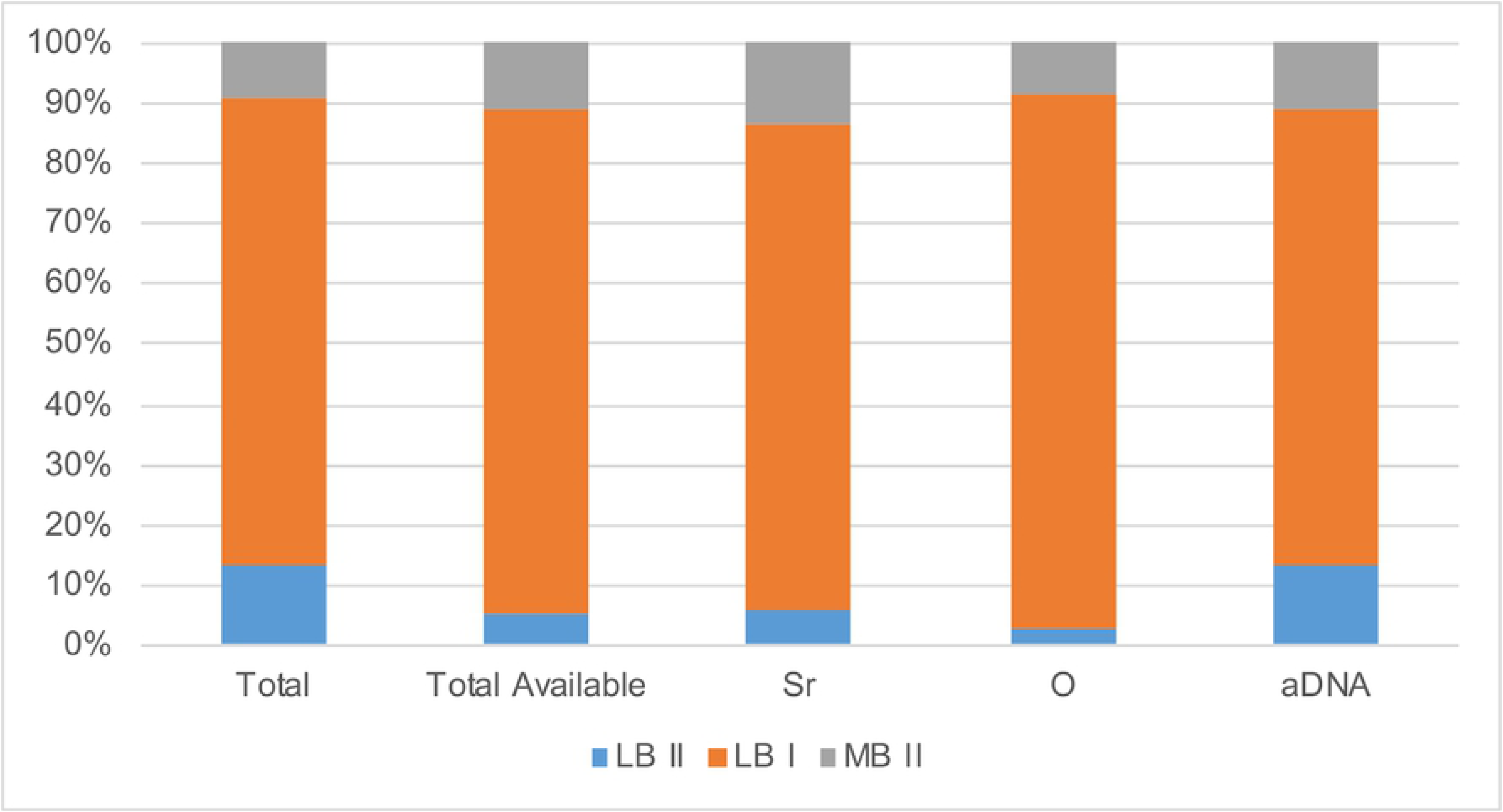
Relative dating based on stratigraphy and context of the total assemblage available for sampling (i.e., excavated by Yener), as well as those sampled for each analysis presented.

**Fig 6.**
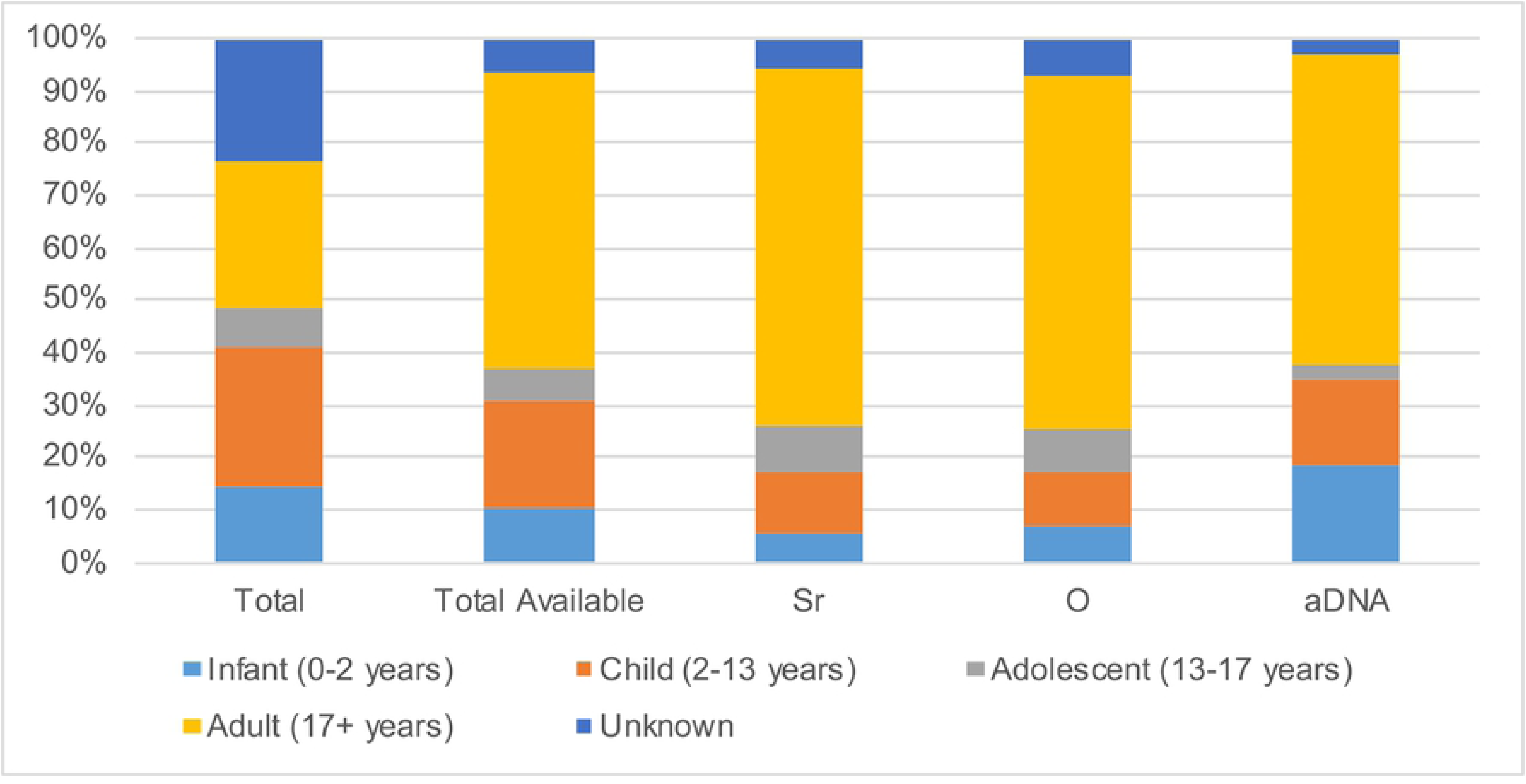
Ages of the total assemblage available for sampling (i.e., excavated by Yener), as well as those sampled for each analysis presented.

However, the excavated burials certainly do not represent the total population who lived and died at the city over the course of its history. It is possible that large swathes of individuals who lived at the site are currently missing from view due to their graves either not having been preserved due to taphonomic processes, not (yet?) having been recovered, or perhaps being archaeologically invisible, due to practices such as off-site burial.

### Isotopic analysis background

The key principle in applying δ^18^O and ^87^Sr/ ^86^Sr values to the study of past mobility is a comparison between the isotopic composition in the tooth enamel of excavated individuals and the hydrologically and biologically available signatures at the same place. If a person spent their childhood prior to the completion of enamel formation of sampled permanent teeth at a different place than their adulthood (typically taken to be represented by the place where the individual was buried), this should result in a mismatch between the δ^18^O and/or ^87^Sr/ ^86^Sr values in their teeth versus the environment, provided the bioavailable isotopic signatures of both places differ from one another [35, 89, 93–103].

Stable oxygen isotopes (δ^18^O) of human tooth enamel are mainly derived from drinking water [94, 97, 99–101] which, in turn, is determined by the interaction of several factors, most importantly, elevation, temperature, humidity, and distance from the sea [93, 95, 96]. In the Amuq Valley, δ^18^O values of modern precipitation average between -7‰ and -6‰ (Fig 7) [104–107], which is also consistent with measured Orontes water values from Syria [108–110]. However, climate change could have altered the bioavailable oxygen over time, and therefore intra-population analysis is generally the preferred method of evaluating δ^18^O results [111], as well as comparisons to faunal samples.

**Fig 7.**
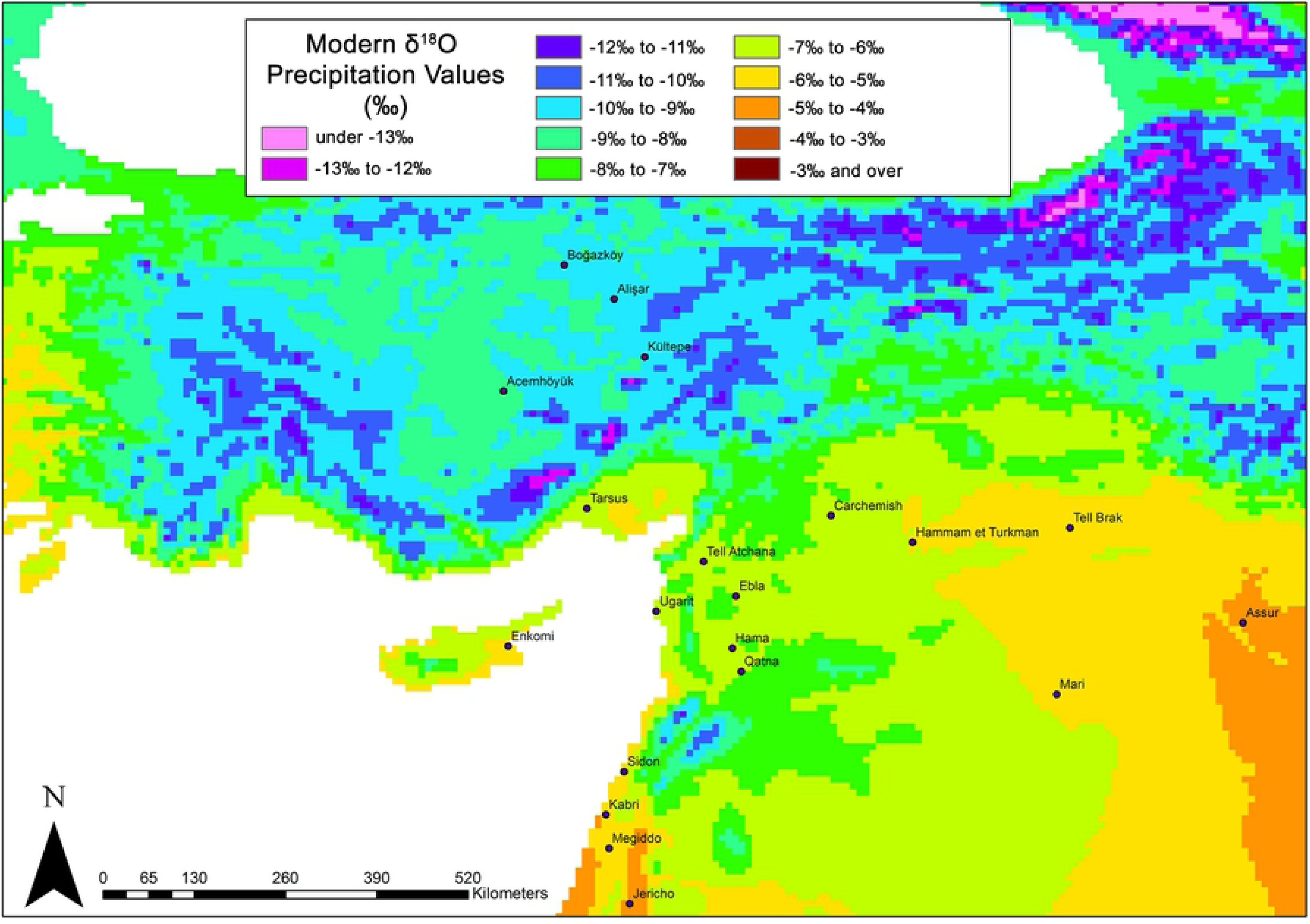
Mean annual δ^18^O values for modern precipitation in the regions surrounding Tell Atchana. Isotopic data from OIPC [104–106].

Strontium in the human body, on the other hand, is incorporated via both food and water, with the biologically available ^87^Sr/ ^86^Sr composition at a location depending mainly on the underlying geological formations. ^87^Sr forms during the radioactive decay of ^87^Rb; therefore, while the amount of ^86^Sr in each rock is stable, the amount of ^87^Sr varies depending on the type of rock (which determines the initial quantity of ^87^Rb and total Sr) and the rock’s age. Weathering processes wash the strontium into soils and runoff water, where it is taken up by plants and then passed on to humans and animals alike, being incorporated into skeletal tissue and teeth during mineralization, as a substitute for calcium, without significant isotopic fractionation [35, 89, 98].

A knowledge of local geology is therefore crucial in order to establish a baseline for strontium isotopic studies. The surface of the Amuq Plain itself is made up mainly of alluvial sediments from the three major rivers (the Orontes, the Kara Su, and Afrin) and eroded material from the highlands surrounding it [112] (Fig 8). The highlands to the south of the valley, which are part of the Arabian Platform [113], are made up of mostly limestone and other carbonate rocks of relatively young age (mainly Miocene and Eocene formations; ^87^Sr/^86^Sr values typically of 0.707-0.709 [114]). There are areas of basalt bedrock in some parts of the Kurt Mountains to the south, which are mostly from the Miocene and Eocene [112, 115], and these can be expected to have somewhat lower ^87^Sr/^86^Sr values (in the range of 0.703-0.705 [116, 117]). Basalt of a somewhat later age, from the Pliocene, can also be found in the northeast of the plain [113, 118, 119], and these areas may be expected to have roughly similar ^87^Sr/^86^Sr values.

**Fig 8.**
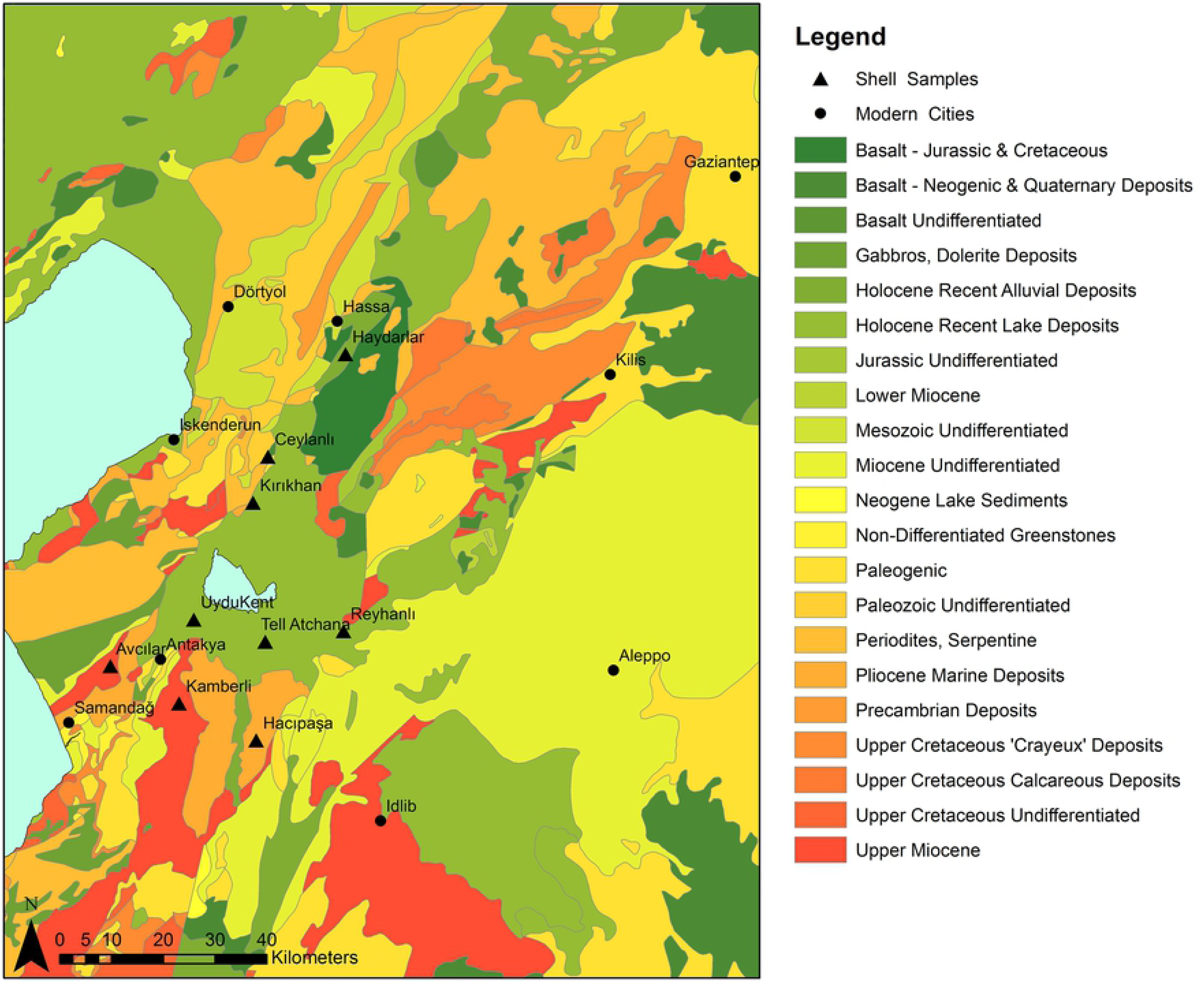
Geological map of the Amuq Valley and surrounding regions, with modern snail sample locations marked. Data courtesy of the Amuq Valley Regional Project.

The Amanus Mountains are much more geologically complex and consist mostly of formations of ultrabasic (or ultramafic) igneous (especially in the southern reaches), metamorphic, and sedimentary rock (particularly in the northern reaches) of more widely varying ages, with some formed as early as the (Pre)Cambrian [112, 115, 118, 120–122], including ophiolites, limestones, gabbro, and basalts, the majority of which are Mesozoic and later in age [118, 123, 124]. The ^87^Sr/^86^Sr values of ophiolites in the Kızıldağ area have been measured as 0.705 [125], and the gabbro fields in the same region can be expected to have similarly low values, comparable to basalt. The southwestern areas of the Amanus range, however, in the area of the Hatay Graben, are composed mainly of carbonates with ^87^Sr/^86^Sr values measured in the range of 0.7088-0.7090 [126]. Further north in the Amanus range, the clastic and carbonate formations are generally older (dating from the Paleozoic and Mesozoic eras) [127] and can therefore be expected to yield higher ^87^Sr/^86^Sr values compared to similar formations on the Arabian Platform to the south.

A strontium isotopes pilot study was conducted by D. Meiggs [128] at Tell Atchana which focused mainly on archaeological faunal and modern environmental samples, although three human samples were also included. The modern environmental samples included both snail shells (six samples) and plants (six samples) collected from various locations around the valley, including one snail shell directly from Tell Atchana (sample AK01), and several of the unanalyzed shells collected during this project were used in the current study in order to compare the two sets of results (for further details, see S2 Text). The ^87^Sr/^86^Sr values of modern samples analyzed by Meiggs ranged from 0.707851-0.714678 with a mean of 0.708998 [128], with the widest variation in ^87^Sr/^86^Sr values found in the samples from the Amanus Mountains (0.707851-0.714678), consistent with the varied geology encountered here. The samples from the alluvial plains of the valley floor showed comparatively lower ^87^Sr/^86^Sr variation (from 0.707942-0.708330), irrespective of if they originated from the northern or southern part of the valley [128]. The snail shell from the tell (AK01: 0.708550) had a slightly higher strontium ratio than those from the plain floor, but was within the range of the ancient faunal samples. The archaeological faunal samples analyzed by Meiggs consisted of teeth from eight ovicaprines and two deer. They provide a much smaller range of ^87^Sr/^86^Sr results, from 0.708196-0.70875, with a mean of 0.708396 [128]. This dataset therefore provides a local ^87^Sr/^86^Sr range (±2 standard deviations from the mean) of 0.708073-0.708718 that likely indicates where strontium signatures of individuals growing up at Alalakh or in its direct vicinity could be expected to fall, although herding practices may have included the use of pastures located on different soils than those used for crop cultivation. In this case, the available ancient faunal samples may not provide a sufficient representation of variation expected in humans.

These considerations show that it is crucial to evaluate where the majority of the food consumed by the individuals under study came from: only if the bulk of the diet was produced locally – i.e., at or in the vicinity of the site where an individual lived – will the strontium isotopic signature allow conclusions about the place of residency, and therefore questions of dietary make-up and catchment must be taken into account [35, 129, 130]. The archaeobotanical evidence of Alalakh is dominated by free threshing wheat (*Triticum aestivum/durum*) and barley (*hordeum vulgare*) [131], although pulses also make up significant portions of the assemblage in certain contexts, including lentils (*Lens culinaris*), fava beans (*Vicia faba*), and chick pea (*Cicer arietinum*) [132]. The Amuq Plain is well-situated for growing these plants, as it lies within the Mediterranean climate region, and an annual mean of 500-700 mm of precipitation, combined with seasonal flooding, allows for rain-fed cereal agriculture on a large scale [115, 133]. The faunal remains recovered from the site consist primarily of domesticates, namely a mix of cattle, sheep/goat, and pig, while wild taxa make up a considerably smaller percentage in most strata [134], although reaching levels as high as 31% in some contexts [135]. This means most animals that were consumed were not roaming free within the Amuq Valley but were managed by people. Occasional consumption of freshwater fish and shellfish occurred, based on their presence in the zooarchaeological assemblage, but not in significant quantities [135]. This suggests that the majority of the daily dietary input of Alalakh’s citizens could have been produced locally.

However, not all of the food present at Alalakh was produced in the immediate vicinity of the site: texts from the palace archives in Periods 7 (MB II) and 4 (LB I) describe regular shipments of food (including barley, emmer wheat, vetches, animal fodder, oil, beer, wine, and birdseed [e.g., texts AlT 236-308b, 320-328]) from Alalakh’s vassal territories [55], and this non-local food, depending on where it was from and the bioavailable strontium of those areas, could have affected the strontium values of individuals who ate it. Most of the identified places where foodstuffs and animals were delivered from were within the control of Alalakh and seem to have been from the Amuq Valley and its immediate environs, although Emar also delivered grain and sheep during Alalakh’s sovereignty over that city in Period 7 [53], demonstrating that not all of the cities under Alalakh’s sway were within the valley. It is unclear what the ultimate destination(s) of these received foodstuffs were – whether they were consumed by the palace denizens, redistributed to palace dependents, given as payment for services or against palace debts, or sold to other residents of Alalakh – but if certain portions of the population were consuming them in large proportions, this has the potential to change their ^87^Sr/^86^Sr ratios and to artificially inflate the numbers of non-locals identified.

### DNA analysis background

The investigation and interpretation of genetic patterns of diversity between humans and groups of humans, usually referred to as populations, is one objective of the field of population genetics. One major factor that shapes genetic variation between populations is geographic distance, as groups living closer to each other are naturally more likely to admix – meaning that individuals are more likely to procreate – than groups living farther apart [136, 137]. Another major factor involved in shaping genetic variation is time, due to continuous human mobility on different scales. The interpretive power of a single-site study such as the current one strongly depends on the availability of already published data of coeval and earlier periods from the Amuq Valley and the wider Near East and Anatolia in general (see below). Furthermore, to securely detect changes in the local gene pool and identify outlier individuals or even different genetic clusters within one place, data from many individuals and archaeological contexts are necessary.

One major difficulty in genetic studies in connection with the identification of genetic outliers at a place concerns the dating of this signal. Often, when an outlier is identified, it is rather difficult to establish whether the sampled individual itself or his/her ancestors immigrated. The combination of aDNA analysis with strontium and oxygen isotope analysis of the same individual is one way to resolve this issue, as migrants in the first generation can be identified, given the isotope signal in their teeth deviates from local baselines. On the other hand, the signal for a first-generation immigrant in the isotopic data can potentially be more closely refined by the aDNA data, due to the general geographic patterning of population genomic data. If an individual identified as a first-generation immigrant by isotopic analysis looks genetically very much like the other individuals at the site, it is likely that we are dealing with either regional/short distance migration or long-distance backwards migration.

In addition to the analysis of genetic ancestry between individuals within one site and between populations, aDNA analysis allows the detection of biological relationships amongst individuals. In some cases, pedigrees can be reconstructed from these [45, 138] which, from an archaeological point of view, can shed light on particular pedigree-related dynamics and practices at a site.

The earliest, and to date only, glimpse into the genetic makeup of the inhabitants of the Amuq Valley prior to Alalakh comes from six samples from Tell Kurdu, five of which date to the Early Chalcolithic between 5750-5600 BC, and one of which is dated to the Middle Chalcolithic, 5005-4849 cal BC (2σ) [46]. Skourtanioti et al. [46] showed, with three different analyses (PCA, *f_4_*-statistics, and *qpAdm*), that the Chalcolithic samples from Tell Kurdu harbor ancestries related primarily to western Anatolia and secondarily to the Caucasus/Iran and the Southern Levant, suggesting a gradient of ancestries with geographical characteristics already in place during that time in the Amuq Valley [46]. However, the samples from the MBA and LBA from Alalakh draw a genetic picture of the Amuq that is considerably changed: roughly 3000 years after the last individual from Tell Kurdu, the individuals from Alalakh, along with individuals from EBA and MBA Ebla in northwestern Syria, are part of the same PC1-PC2 space with Late Chalcolithic-Bronze Age Anatolians. They are, compared to samples from Barcın in western Anatolia and Tell Kurdu, all shifted upwards on the PC2 towards samples of Caucasus and Zagros/Iranian origin (Fig 9) [46]. This shift in ancestry was formally tested with *f_4_-*statistics of the format *f_4_*(Mbuti, test; Barcın_N/TellKurdu_EC, X), which revealed that all the Late Chalcolithic-LBA populations from Anatolia and the northern Levant (X, i.e. Ebla and Alalakh) are more closely related to Iranian Neolithic individuals and/or Caucasus Hunter Gatherer individuals (*test*) than are the earlier Tell Kurdu and Barcın individuals [46]. A similar genetic shift towards Iranian/Caucasus-related populations was detected for contemporary Southern Levant [139–141]. This means that in the period between 5000–2000 BC, gene flow from populations harboring Iranian/Caucasus-like ancestries, which also includes populations that are genetically similar to these but have not yet been sampled, and are thus unknown, affected southern Anatolia and the entire Levant, including the Amuq Valley. It is currently neither possible to pinpoint the exact source population(s) that brought about these changes in the local gene pool nor to propose specific migration events.

**Fig 9.**
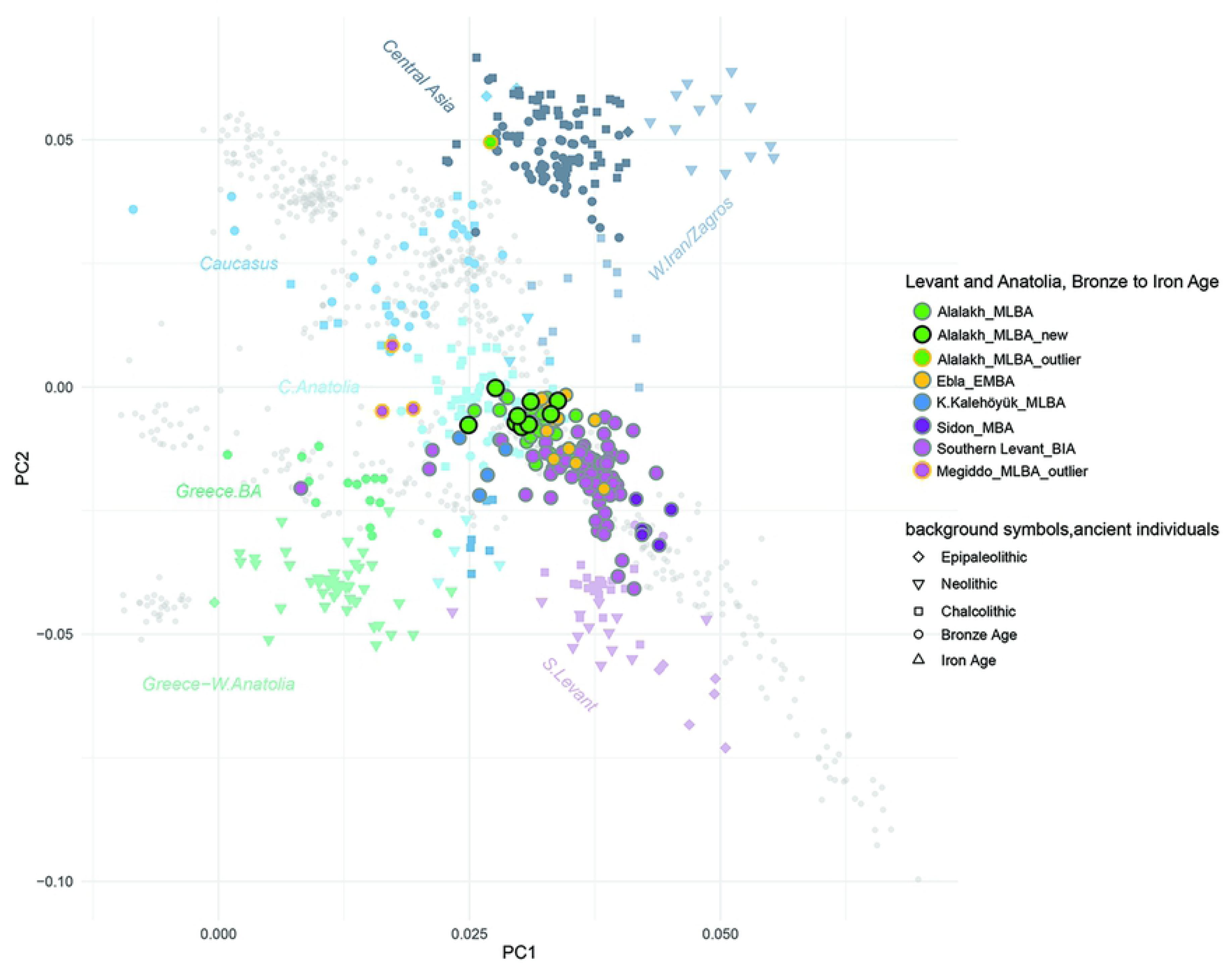
PCA: scatterplot of PC1 and PC2 calculated on West Eurasian populations (Human Origins dataset; grey symbols) using smartpca with projection of ancient individuals (colored symbols).

Four genetic outlier individuals from Bronze Age Levantine contexts, one of them the so-called Well Lady from Alalakh (ALA019) and three from Megiddo (two of which are siblings), are shifted upwards on the PCA, the former towards individuals from Chalcolithic/Bronze Age Iran and Central Asia [46] and the latter to the Chalcolithic/Bronze Age Caucasus. Strontium isotope analysis of the two siblings from Megiddo suggests that both grew up locally [139]. These outlier individuals from Megiddo and Alalakh attest that gene flow from Caucasus/Iran (or genetically similar groups) into the Levant continued throughout the second millennium BC.

### Analytical methods

#### Stable oxygen isotopes

Sampling protocols and analysis procedures for stable oxygen isotope analysis follow those set out in Roberts et al. [142] (see also [143–145]). Teeth were cleaned to remove adhering material using air-abrasion, and a diamond-tipped drill was used to obtain a powder sample. The full length of the buccal surface was abraded in order to capture a representative bulk sample from the maximum period of formation. To remove organic or secondary carbonate contamination, the enamel powder was pre-treated in a wash of 1.5% sodium hypochlorite for 60 minutes; this was followed by three rinses in purified H_2_O and centrifuging, before 0.1 M acetic acid was added for 10 minutes. Samples were then rinsed again three times with milliQ H_2_O and freeze dried for 4 hours. Enamel powder was weighed out into 12 ml borosilicate glass vials and sealed with rubber septa. The vials were flush filled with helium at 100 ml/min for 10 minutes. After reaction with 100% phosphoric acid, the CO_2_ of the sample was analyzed using a Thermo Gas Bench 2 connected to a Thermo Delta V Advantage Mass Spectrometer at the Stable Isotope Laboratory, Department of Archaeology, MPI-SHH. δ^13^C and δ^18^O values were calibrated against International Standards (IAEA NBS 18 : δ^13^C -5.014 ± 0.032‰; δ^18^O - 23.2±0.1‰, IAEA-603 [δ^13^C = +2.46±0.01‰, δ^18^O -2.37±0.04‰]; IAEA–CO–8 [δ^13^C - 5.764±0.032‰, δ^18^O -22.7±0.2‰,]; USGS44 [δ^13^C = −42.1‰,]). Repeated analysis of MERCK standards suggests that machine measurement error is ca. +/- 0.1‰ for δ^13^C and +/- 0.1‰ for δ^18^O. Overall measurement precision was determined through the measurement of repeat extracts from a bovid tooth enamel standard (n = 20, ±0.2‰ for δ^13^C and ±0.3‰ for δ^18^O).

#### Strontium isotopes

Sampling protocols and analytical procedures for strontium follow those set out in Copeland et al. [146]. Enamel powder was obtained with a diamond-tipped drill along the full length of the buccal surface after cleaning with air-abrasion. Up to 4 mg of enamel powder was digested in 2 ml of 65% HNO_3_ in a closed Teflon beaker placed on a hotplate for an hour at 140°C, followed by dry down and re-dissolving in 1.5 ml of 2 M HNO_3_ for strontium separation chemistry, which followed Pin et al. [147]. The separated strontium fraction was dried down and dissolved in 2 ml 0.2% HNO_3_ before dilution to 200 ppb Sr concentrations for analysis using a Nu Instruments NuPlasma High Resolution Multi Collector Inductively Coupled Plasma-Mass Spectrometry (HR-MC-ICP-MS) at the Department of Geological Sciences, University of Cape Town. Analyses were controlled by reference to bracketing analyses of NIST SRM987, using a ^87^Sr/^86^Sr reference value of 0.710255. Data were corrected for rubidium interference at 87 amu using the measured ^85^Rb signal and the natural ^85^Rb/^87^Rb ratio. Instrumental mass fractionation was corrected using the measured ^86^Sr/^88^Sr ratio and the exponential law, along with a true ^86^Sr/^88^Sr value of 0.1194. Results for repeat analyses of an in-house carbonate reference material processed and measured as unknown with the batches (^87^Sr/^86^Sr = 0.708909; 2 sigma 0.000040; n = 7) are in agreement with long-term results for this in-house reference material (^87^Sr/^86^Sr = 0.708911; 2 sigma = 0.000040; n = 414).

#### aDNA

DNA data production of all nine newly sampled individuals in this study took place in the dedicated aDNA facility of the MPI-SHH in Jena, Germany. Sampling targeted the inner-ear part of the petrous bone [87]. DNA extraction and double-stranded genomic libraries were prepared for four samples (ALA118, ALA120, ALA123, and ALA124) according to the MPI-SHH Archaeogenetics protocols for Ancient DNA Extraction from Skeletal Material, and Non-UDG treated double-stranded ancient DNA library preparation for Illumina sequencing, both archived and accessible at dx.doi.org/10.17504/protocols.io.baksicwe and dx.doi.org/10.17504/protocols.io.bakricv6, respectively. The library preparation protocol was modified with the introduction of partial Uracil DNA Glycosylase (UDG) treatment prior to the blunt-end repair, according to Rohland et al. [148]. Dual-indexed adaptors were prepared according to the archived MPI-SHH Archaeogenetics protocol accessible at dx.doi.org/10.17504/protocols.io.bem5jc86.

For the remaining five samples (ALA130, ALA131, ALA135, ALA136, and ALA138), DNA extraction was performed according to Rohland et al. [149], and single-stranded libraries (no UDG treatment) were prepared according to Gansauge et al. [150], both protocols using an automated liquid-handling system. All libraries were first shotgun sequenced (∼5M reads) in a sequencing Illumina HiSeq4000 platform. Raw FastQC sequence data were processed through EAGER [151] for removal of adapters (AdapterRemoval [v2.2.0]) [152], read length filtering (>30b), mapping against hs37d5 sequence reference (BWA [v0.7.12]) [153], q30 quality filter, removal of PCR duplicates (dedup [v0.12.2]) [151], and DNA damage estimation (mapdamage [v2.0.6]) [154]. Two main characteristics of the sequenced reads were considered in order to select positive libraries for submission to an in-solution hybridization enrichment that targets 1,233,013 genome-wide and ancestry-informative single nucleotide polymorphisms (SNPs; “1240K SNP capture”) [155]. The first one is the proportion of DNA damage at the end of the reads (> ∼5% C-T/ G-A substitution at terminal 5’ and 3’ base, depending on the UDG treatment of the library), and the second one is the content of endogenous DNA > 0.1%, the latter calculated as the portion of reads mapped against the hs37d5 reference over the total amount of sequenced reads after the length filtering. “Captured” libraries were sequenced at the order of ≥20M reads and the raw FastQC sequence data were processed through EAGER. We created masked versions of the bam files using trimBam (https://genome.sph.umich.edu/wiki/BamUtil:_trimBam) by masking the read positions with high damage frequency, that is the terminal 2 and 10 bases for the partially UDG-treated double-stranded libraries, and single-stranded (no UDG) libraries, respectively. We used “samtools depth” from the samtools (v1.3) [156] on the masked bam files providing the bed file with the 1240K SNPs to calculate the coverage on X, Y, and autosomal chromosomes. X and Y coverage were normalized by the autosomal coverage (X-rate and Y-rate respectively), and females without contamination were determined by X-rate ≈ 1 and Y-rate ≈ 0, whereas males without contamination were determined by both rates ≈ 0.5. We used the original bam files in order to estimate mitochondrial contamination with Schmutzi [157] and the nuclear contamination on males with ANGSD (method 1) [158].

We called genotypes with the tool pileupCaller (https://github.com/stschiff/sequenceTools/tree/master/src/SequenceTools) according to the Affymetrix Human Origins panel (∼600K SNPs) [159, 160] and the 1240K panel [155]. We used the option randomHaploid which randomly draws one read at every SNP position. We performed the random calling both on the original and the masked bam files of each double-stranded library, and, for the final genotypes, we kept transitions from the masked and from the original bam files. We used only the original bam files from the single-stranded libraries, and we applied the singleStrandMode option that removes reads with post-mortem damage based on their alignment on the forward or the reverse strand of the human reference genome. We report information about library processing, genetic sex, damage patterns, SNP coverage, and contamination in Table 5.

**Table 5.**
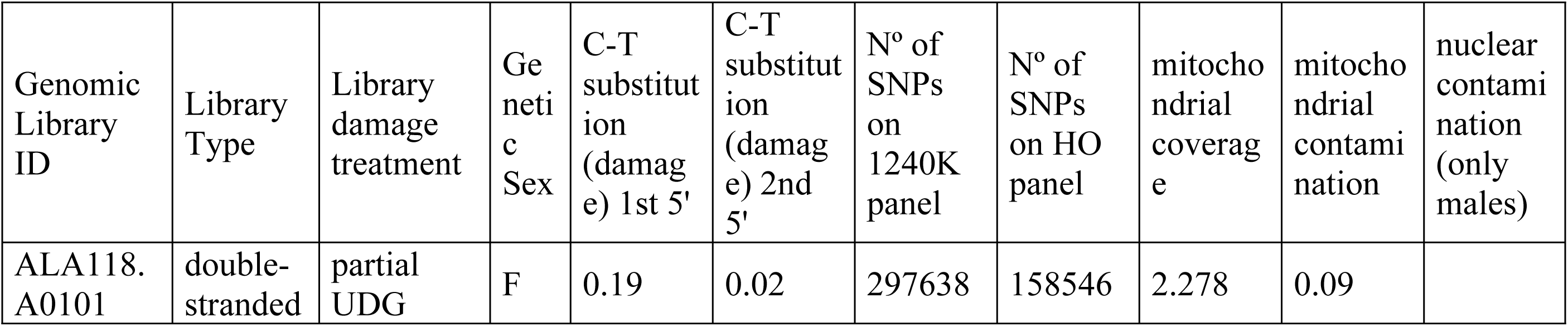
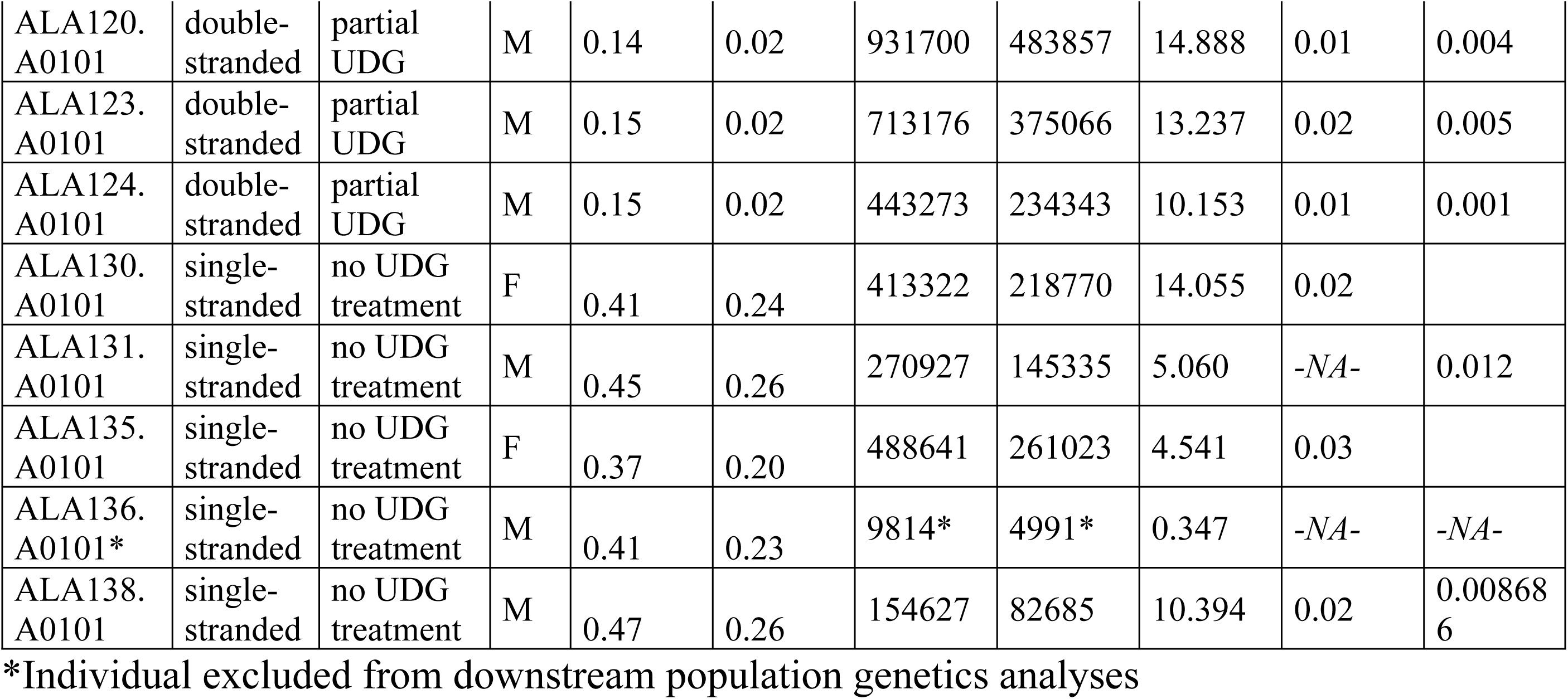
aDNA data of new individuals from Alalakh published in this study.

| Genomic Library ID | Library Type | Library damage treatment | Genetic Sex | C-T substitution (damage) 1st 5' | C-T substitution (damage) 2nd 5' | N° of SNPs on 1240K panel | N° of SNPs on HO panel | mitochondrial coverage | mitochondrial contamination | nuclear contamination (only males) |
| --- | --- | --- | --- | --- | --- | --- | --- | --- | --- | --- |
| ALA118. A0101 | double-stranded | partial UDG | F | 0.19 | 0.02 | 297638 | 158546 | 2.278 | 0.09 |  |
| ALA120.<br>A0101 | double-stranded | partial<br>UDG | M | 0.14 | 0.02 | 931700 | 483857 | 14.888 | 0.01 | 0.004 |
| ALA123.<br>A0101 | double-stranded | partial<br>UDG | M | 0.15 | 0.02 | 713176 | 375066 | 13.237 | 0.02 | 0.005 |
| ALA124.<br>A0101 | double-stranded | partial<br>UDG | M | 0.15 | 0.02 | 443273 | 234343 | 10.153 | 0.01 | 0.001 |
| ALA130.<br>A0101 | single-stranded | no UDG<br>treatment | F | 0.41 | 0.24 | 413322 | 218770 | 14.055 | 0.02 |  |
| ALA131.<br>A0101 | single-stranded | no UDG<br>treatment | M | 0.45 | 0.26 | 270927 | 145335 | 5.060 | -NA- | 0.012 |
| ALA135.<br>A0101 | single-stranded | no UDG<br>treatment | F | 0.37 | 0.20 | 488641 | 261023 | 4.541 | 0.03 |  |
| ALA136.<br>A0101* | single-stranded | no UDG<br>treatment | M | 0.41 | 0.23 | 9814* | 4991* | 0.347 | -NA- | -NA- |
| ALA138.<br>A0101 | single-stranded | no UDG<br>treatment | M | 0.47 | 0.26 | 154627 | 82685 | 10.394 | 0.02 | 0.00868<br>6 |

\*Individual excluded from downstream population genetics analyses

Due to the low coverage of ALA136 (<1% of 1240K sites), we excluded this individual from downstream population genetics analyses. We combined the data from the remaining individuals with previously published ancient and modern individuals [46, 85, 141, 159, 161–190]. For readability, we kept most of the group labels used in Skourtanioti et al. [46], most importantly “Alalakh_MLBA”, “ALA019” (genetic outlier) (n = 1), “Ebla_EMBA” (n = 9), “K.Kalehöyük_MLBA” (Kaman-Kalehöyük, n = 5), and “TellKurdu_EC” (n = 5), but dubbed individuals from Sidon with the label “Sidon_MBA” (instead of “Levant_MBA”; n = 5). We performed principal component analysis on a subset of western Eurasian populations of the Human Origins Dataset using smartpca program of EIGENSOFT (v6.01) [191, 192] (default parameters and options lsqproject: YES, numoutlieriter:0) (see Fig 9).

We assessed the degree of genetic relationship among Alalakh_MLBA individuals (n = 34 after quality filtering) by applying and comparing two different methods: READ [193] and *lcMLkin* [194]. Read is a software that can estimate up to second degree relationships from low-coverage genomes by calculating the proportion of non-matching alleles for a pair of individuals (PO) in non-overlapping windows of 1 Mbps. P0 was normalized with the median of P0 from all pairs – assuming that most pairs are unrelated – in order to reduce the effects of SNP ascertainment, within-population diversity, and marker density.

*LcMLkin* uses a Maximum Likelihood framework on genotype likelihoods from low-coverage DNA sequencing data and infers k0, k1, and k2, the probabilities that a pair of individuals share, respectively, zero, one or two alleles identical-by-descent (IBD), as well as the overall coefficient of relatedness (*r*). Two useful aspects of this method are that it serves for distinguishing between parent-offspring (k0=0) and siblings (k0≥0, depending on recombination rate) and can infer relatedness down to the 5^th^ degree. However, a discrepancy from the expected k0, k1, k2, and *r* values can occur under scenarios of recent admixture, inbreeding, contamination, and low-quality data. We run *lcMLkin* on the masked bam files with the options -l phred and -g best.

We used *qpWave* and *qpAdm* programs from ADMIXTOOLS [160] for modelling of ancestry proportions, using the following set of Right populations (also named outgroups or references): Mbuti.DG, Ami.DG, Onge.DG, Mixe.DG, Kostenki14, EHG, Villabruna, Levant_EP, and Barcın_N. These programs compute a matrix of *f_4_*-statistics for the Right and Left (targets for *qpWave* and target and sources for *qpAdm*) populations in the form of *F*_*ij*_=*F*_4_(*L*_1_,*L*_*j*_;*R*_1_,*R*_*j*_). Then, with a likelihood ratio test, the null model is compared against the full-rank model in which all columns of the matrix are independent. In the latter model, the *n* Left populations relate with the references through *n* waves of ancestry, which for *qpAdm*, implies that the target cannot be explained as a combination of the selected source populations (null model). Depending on the chosen cutoff, a tested null model with p-value ≤0.01 or ≤0.05 and/or infeasible admixture coefficients (outside 0-1 range) is rejected. For this group-based analysis, we kept only individuals who are not genetically related.

## Results

### Results of oxygen isotope analysis

The 77 individuals analyzed yielded a mean δ^18^O of -5.2±0.9‰ and a range of 4.1‰ (from -7.3±0.1‰ to -3.2±0.1‰; Fig 10, Table 4, S1 Table), with values clustering mainly between -6.0‰ and -4.0‰. There are no statistically significant differences identified by one-way ANOVA test among the population according to age, sex, burial type/location/goods, archaeological period, etc.

**Fig 10.**
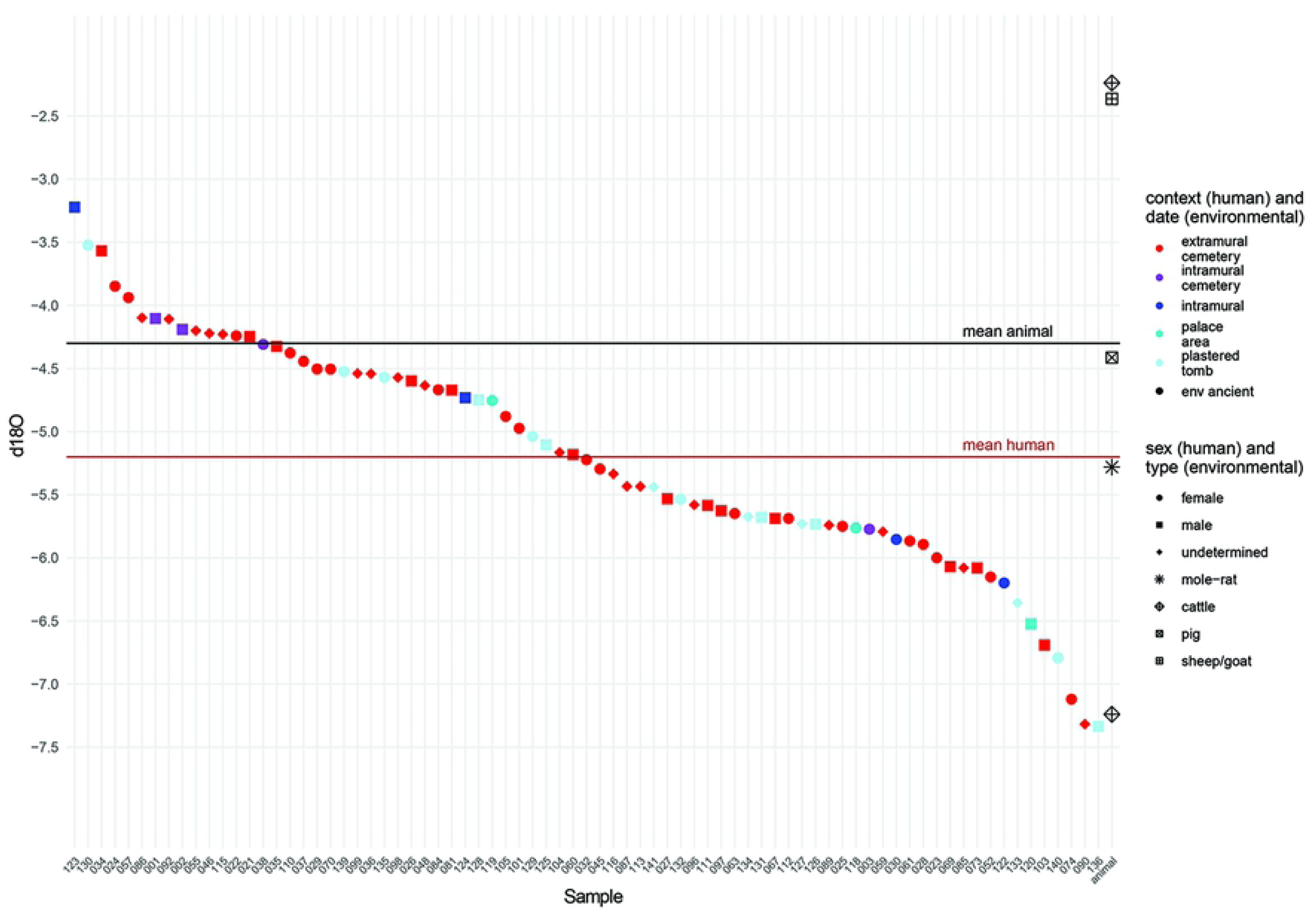
All δ^18^O results.

Following recent suggestions that in-group statistical methods to identify outliers is a more reliable way of identifying non-locals within sets of δ^18^O values than ranges of variation, which have been shown to be ca. 3‰ within a population [40, 111], there are no clear statistical outliers among the Tell Atchana dataset. The five archaeological faunal samples (all from domestic animals; see Table 3) have a higher mean of -4.3±2.1‰ and a wider range of 5‰ (from -2.2±0.05‰ to -7.2±0.05‰). This is due to two particularly high results from AT 0263 and AT 3064, a cattle and an unidentified sheep/goat, respectively. Nevertheless, the results of the humans and fauna are broadly compatible.

### Results of strontium isotope analysis

Every strontium isotope study is faced with the challenge of how best to establish the local bioavailable ^87^Sr/ ^86^Sr range at the site under study. While two standard deviations from the mean have become common practice to set an objective cut-off to distinguish locals from non-locals [98], the material on which to base this mean is debated and varies between different studies. In this study, we used a mixed approach between ancient (snail shells, rodent teeth, sheep/goat teeth, and deer teeth) and modern faunal samples (snail shells) to establish (1) a local range for Alalakh and (2) a local range for the Amuq Valley in general, in order to be able to distinguish between those human individuals that grew up at Alalakh (locals), those who came to the site from within the Amuq (micro-regional migration), and those originating from places outside the Amuq Valley (non-locals: migration over longer distances).

To estimate the typical local ^87^Sr/ ^86^Sr signature for humans at Alalakh, we measured, in addition to the existing samples from sheep/goat and deer teeth [128], ^87^Sr/ ^86^Sr ratios of five land snail shells and tooth enamel from two rodents from well-stratified archaeological contexts (see Table 3). As opposed to domesticates, rodents and snails are not managed by humans, and they obtain their food from within a small radius that should be representative for the strontium ratios available directly at the tell [195]. The snails and rodents offer a means to control whether the ovicaprines were grazing on pastures around Tell Atchana itself within the Amuq Valley, where the bulk of the humans’ plant diet was likely produced, or whether the pastures were located on different geologies (in more mountainous areas on the fringes of the Amuq Valley). The ^87^Sr/ ^86^Sr values of the ancient snail shells and rodents clustered closely together between 0.708111 and 0.708544 (Fig 11) and largely overlapped with the ^87^Sr/ ^86^Sr ratios of the ovicaprines and deer from Meiggs’ study [128], but, as expected, considering the differences in radius of movement, the ratios of the ovicaprines and deer have a wider range. Therefore, we can report positive results for the use of land snail shells as material to obtain bioavailable strontium signatures at Tell Atchana, contributing to a lively discussion in the literature where they have been used with varying success rates [195–201]. The snails and rodents confirm that herding practices of the ovicaprines mostly included pastures in the environs of Alalakh. Thus, the combination of the ovicaprines and deer, together with the two rodents, likely indicates the most relevant local range to represent locality in humans at Alalakh, returning a local range as two standard deviations from the mean (0.708401) of 0.708085-0.708717 (Table 6). By excluding the five snail shells from this calculation, we avoid a potential bias stemming from the snails’ fixation to a very small radius on the tell that may be less representative for humans.

**Fig 11.**
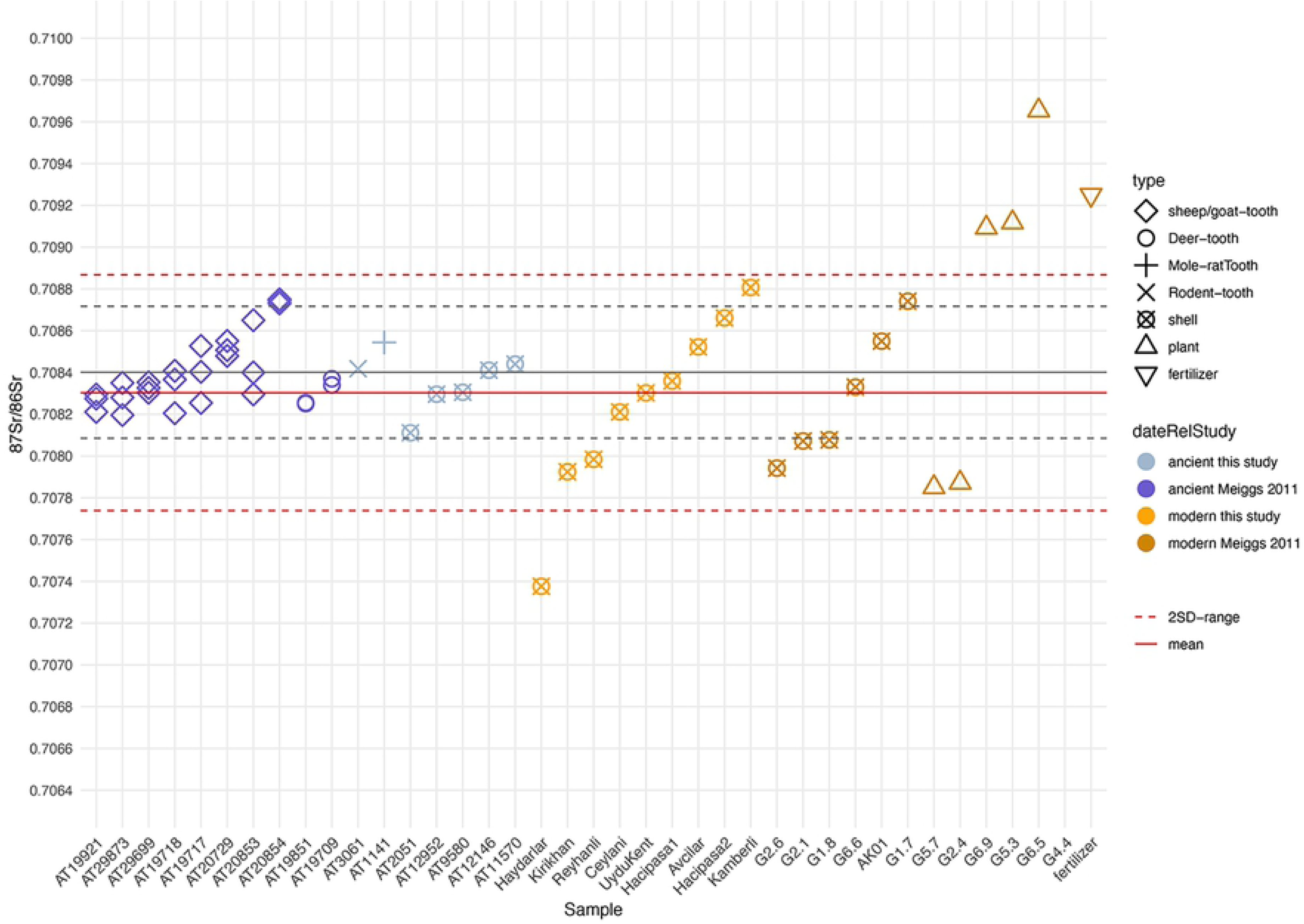
^87^Sr/^86^Sr ratios of snail, plant, fertilizer, and animal samples from this study and Meiggs [128]; continuous lines: mean; dotted lines: ±2 SD from mean; dark grey lines: local range for Alalakh, calculated from sheep/goat, deer, and rodents’ teeth; dark red lines: local range for the Amuq Valley, calculated from modern and ancient snail shells of both studies, using the mean of the five ancient snails from Alalakh as representative for this location, and excluding the modern samples AK01 and the outlier from Haydarlar. Note: sample G4.4 falls outside the ranges plotted in this graph and therefore appears blank.

**Table 6.** Comparison between possible local ranges.

| Description | Mean | Local Range (+/-2 SD) | Comments |
| --- | --- | --- | --- |
| All ancient environmental samples from Alalakh (sheep/goat and deer teeth, n = 28) | 0.708396 | 0.708073-0.708718 | Meiggs' study [128] |
| All ancient environmental samples from Alalakh (5 snails, 2 rodents) | 0.708361 | 0.708104-0.708617 | this study |
| All ancient snail shells from Alalakh (n = 5) | 0.708313 | 0.708081-0.708544 | this study |
| All ancient environmental samples of both studies (n=35) | 0.708389 | 0.708077-0.708700 | Meiggs' study [128] and this study |
| **Ancient samples of sheep/goat, deer and rodents' teeth (n=30)** | **0.708401** | **0.708085-0.708717** | **Local range, Alalakh; Meiggs' study [128] and this study** |
| All modern environmental samples from Amuq Valley (snails and plants, n = 12) | 0.708998 | 0.705400-0.712596 | Meiggs' study [128] |
| Modern environmental samples from Amuq Valley, excluding pine needle outlier (Sample G4.4) | 0.708482 | 0.707331-0.709632 | Meiggs' study [128] |
| All modern snail shells from Amuq Valley (n = 6) | 0.708285 | 0.707716-0.708855 | Meiggs' study [128] |
| All modern snail shells (n = 9) | 0.708238 | 0.707420-0.709057 | including Haydarlar outlier; this study |
| **modern snail shells combined with mean of ancient snails (excluding Haydarlar outlier and AK01 from Alalakh) (n = 14)** | **0.708303** | **0.707739-0.708868** | **Local range, Amuq; Meiggs' study [128] and this study** |
| 6 youngest individuals (0-6 years) from Alalakh | 0.708340 | 0.708136-0.708544 | Humans, this study |

One way to check the accuracy of a local range obtained from the ancient faunal samples is by comparison against the ^87^Sr/ ^86^Sr values of young children: the likelihood of individual mobility in sedentary societies should increase with age, so individuals dying at a young age are more likely to be local [200, 202–204]. All six individuals under the age of seven from Alalakh fall well inside the local range as determined by the archaeological fauna. In general, we believe that the range calculated from ancient faunal samples is representative for locality in humans, although in the case of individuals falling just outside this local range, we need to consider the option that these may only appear as outliers, if they were consuming larger portions of non-local diet as compared to other inhabitants.

The modern snail samples taken from throughout the valley provide the opportunity to compare the ^87^Sr/ ^86^Sr values at the site with those from other locations in the Amuq Valley and serve to calculate a local range for the valley in general. The modern snail shells from our study (n = 9) show a high consistency with the snail shells from Meiggs’ study (n = 6), with samples originating from the same geological units having similar ^87^Sr/ ^86^Sr values across both studies. The plant samples from Meiggs’ study, on the other hand, are generally characterized by either extremely high or low ^87^Sr/ ^86^Sr values that cannot be explained by their location within the geological patchwork of the slopes of the Amanus mountains on the fringes of the Amuq Valley alone (for further discussion see S2 Text). We therefore decided to combine only the snail shells of both studies in our calculations of a local range for the Amuq Valley catchment area. The snail from Haydarlar, with the lowest ^87^Sr/ ^86^Sr value (0.707376) among the modern snails, constitutes an outlier compared to all other modern snails. Haydarlar is located on alluvial deposits of the Kara Su river valley on the northernmost fringes of the Amuq Valley. We conclude that the distinctly low isotopic signature of this snail stems from the basalt shields of Jurassic and Cretaceous age that are located along the slopes of the river valley, so that runoff water from these areas is naturally directed toward the riverbed and therefore impacts these adjacent areas, pulling the snail shell toward a lower ^87^Sr/ ^86^Sr value (see also S3 Text) [195, 200]. This does not mean that the result should be considered incorrect, only that individuals growing up around this location may also have a comparable strontium signature that is distinctly lower than that of individuals from the rest of the Amuq Valley. We therefore excluded the snail shell from all further calculations of a local range for the Amuq Valley. Finally, we excluded the modern snail shell from Alalakh itself (sample AK01) and instead used the mean of the ancient snail shells (n = 5), since we expect these to be a better representative for the local signature directly at the tell. With this method, we obtain a local range for the wider Amuq catchment area, based on 14 distinct data points, of 0.707739-0.708868, and a mean of 0.708303 that we see as best representing the strontium variation within the valley.

Applying the local range for Alalakh (708085-0.708717), out of a total of 53 human individuals, 40 plot within the range of Alalakh and another 8 plot outside the Alalakh range, but within the range for the Amuq Valley (0.707739-0.708868) (Fig 12; see also Table 4). Five individuals can be securely identified as non-locals to both Alalakh and the Amuq, plotting outside both local ranges (ALA110, ALA098, ALA037, ALA004, and ALA033). Nearly 10% of the sampled population (9.3%) is therefore identified as non-local to both Alalakh and the Amuq Valley.

**Fig. 12.**
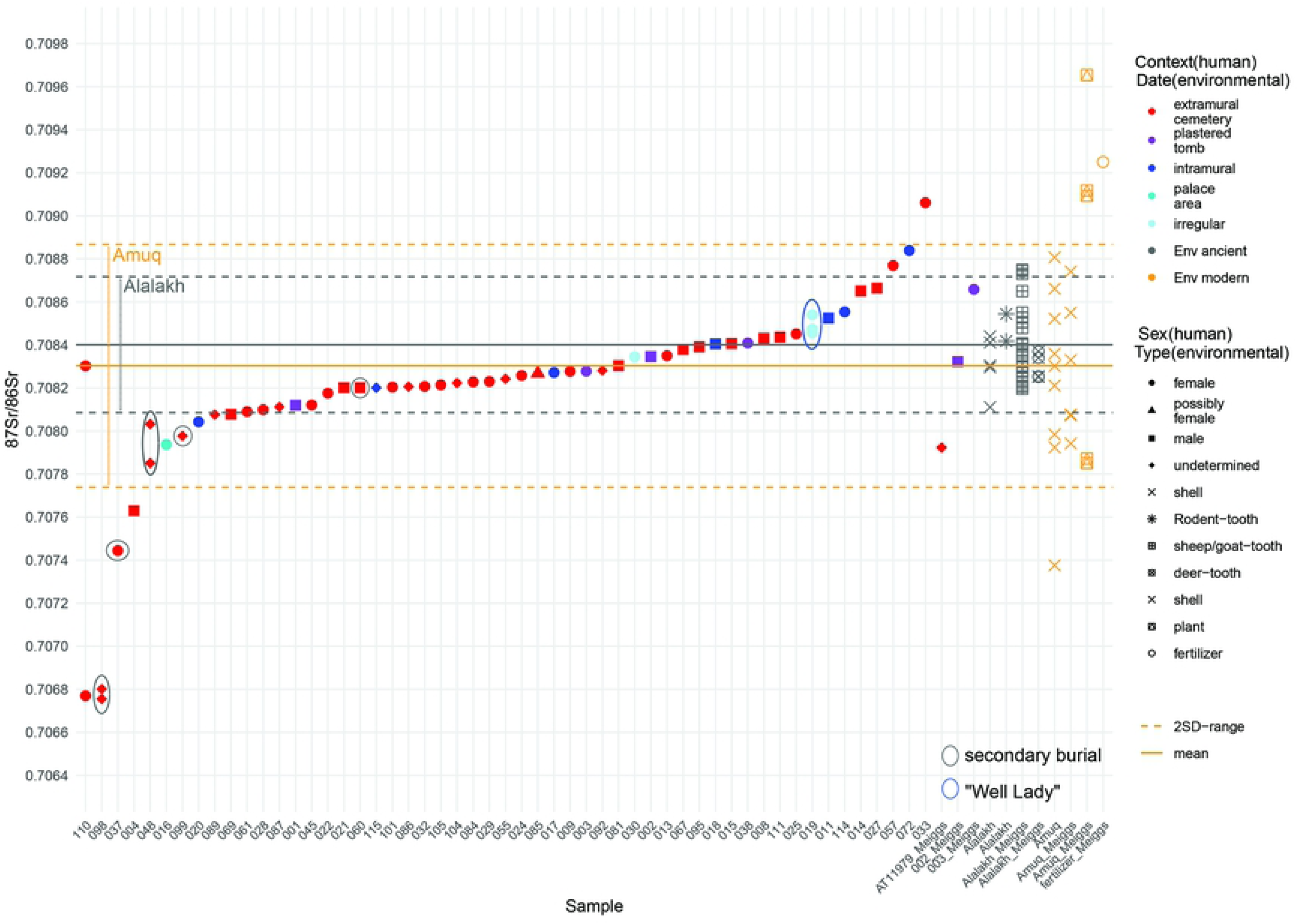
All ^87^Sr/ ^86^Sr results, plotted against local ranges. Black lines: local range calculated from ancient faunal samples from Alalakh; orange lines: local range calculated from modern snails from the Amuq and the mean of ancient environmental samples from Alalakh.

All five non-local individuals were buried in the extramural cemetery, and four of the five are stratigraphically dated to Period 6 (ALA110 is dated to Period 7 and is one of the earliest graves excavated in Area 3; see Table 2). Three are female (two adults – ALA110 and ALA033 – and one of unknown age – ALA037), one is an adult male (ALA004), and one is of unidentified age and sex (ALA098). All three non-locals who have also been analyzed for oxygen isotopes (ALA110, ALA098, and ALA037) fall firmly within the range of local δ^18^O values (see Table 4), indicating that, while they are not from the Amuq Valley, they grew up in areas with similar δ^18^O values. Most interestingly, two of the five non-locals (ALA098 and ALA037) are secondary burials, as are secondarily buried individuals ALA048 and ALA099, two of those who likely came from within the Amuq Valley, rather than Alalakh itself (see Fig 12). In fact, only one of the sampled secondary burials (ALA060) falls within the local range for Alalakh (see Fig 12). In order to explore the timing of the migration of these non-locals, M3s were also analyzed when they were available, which returned a range of resulting patterns (see Fig 12). Like the M2 (0.707851), the M3 of ALA048 (0.708032) still falls within the group identified as local to the Amuq, but substantially closer to the Alalakh range, indicating that the move from within the Amuq to Alalakh may have occurred late during the formation of the M3 (likely during the end of childhood/early adolescence), leading to this mixed signal,. The M3 of ALA110 (0.708303) falls firmly within any local range calculated here and clearly shows that this woman moved to Alalakh in later childhood – i.e., between the formation of M2 and M3. ALA098, however, has similar ^87^Sr/^86^Sr values for both M2 (0.706801) and M3 (0.706755), both of which fall at the lowest end of the results reported here. It therefore appears that this individual spent their entire childhood and youth in another location, moving to Alalakh only in adulthood.

Out of the three human individuals sampled in Meiggs’ study [128], two were analyzed again in this study (002_Meiggs = ALA002 and 003_Meiggs = ALA003). While both samples from individual ALA002 have similar ^87^Sr/ ^86^Sr ratios, ALA003 in Meiggs’ study has a higher ^87^Sr/ ^86^Sr ratio and plots outside the local range calculated here, as does the third human sample from Meiggs’s study (AT 11979). Unfortunately, the teeth sampled by Meiggs were only identified to the level of molars, and, given the discrepancy between the M2 value obtained here and the one published by Meiggs [128], it is likely that the tooth sampled in Meiggs’ study was either an M3 or an M1. In this case, the difference in the ^87^Sr/ ^86^Sr ratio between the two samples from ALA003 would be explained by changes in the origins of food that could ultimately be linked to a change in place of residency during childhood. While a sample from an M1 would mean that ALA003 spent the first years of her life outside of Alalakh, a sample from the M3 would hint at a move away from Alalakh during later childhood/early adolescence and, consequently, a return to Alalakh later.

### Results of aDNA analysis

All individuals sampled from Alalakh, regardless of their context, are very homogeneous from a population genomics perspective, with only one exception (ALA019). As described above (see Table 4), the individuals cover all ages (ca. 40 weeks-75 years at death) and both sexes, as well as all burial contexts available for analysis. It is reasonable to assume, therefore, that the genomic data from Alalakh accurately describes the genetic variation within the bulk of the MBA-LBA population from Alalakh. Published data from other contemporary Levantine and Anatolian sites shows that most individuals cluster relatively close to each other in the PCA on a north-south cline, and their overall genetic differences are small [46, 140, 141, 161], yet detectable. Therefore, with the help of *qpAdm* modeling, we can explore the role of Alalakh as an intermediary on this cline between contemporaneous individuals from sites located to the north in Anatolia and to the south in present day Lebanon [205]. For modeling, we have chosen individuals dating to the MBA and LBA from Kaman-Kalehöyük (n = 5 [161]) as a representative for central Anatolian groups and from Sidon (n = 5 [141]) as a representative for Levantine individuals to the south of Alalakh.

As – at least for the Amuq Valley and the Southern Levant – there was gene flow during and/or after the Chalcolithic period, we tested models that used temporally proximal sources from Anatolia, Iran, the Caucasus, and the Southern Levant (Fig 13 and S2 Table). The results of this modeling show that Alalakh_MLBA (n = 31) can be adequately modeled as a three-way admixture model between an Anatolian (“Büyükkaya_Chl”), a Levantine (“Levant_Chl”), and an Iranian (“Iran_Chl”) source (pval = 0.28), while for Sidon_MLBA, the two-way admixture model of Levant_Chl and Iran_Chl provides the best fit (pval = 0.037). Three-way models fail for Sidon_MLBA (pval < 0.01 or negative coefficients) (see Fig 13). While the same admixture model for Alalakh applies to Ebla_EMBA, with a lower Büyükkaya_EChl ancestry coefficient, nested models such as Iran_Chl (47.2±2.6%) + Levant_Chl (52.8±2.6%) also become adequate (p ≥ 0.5). The fit of simpler models for Ebla might be a result of lower statistical power to distinguish between the model and actual targets, due to their smaller sample size and/or coverage compared to Alalakh.

**Fig 13.**
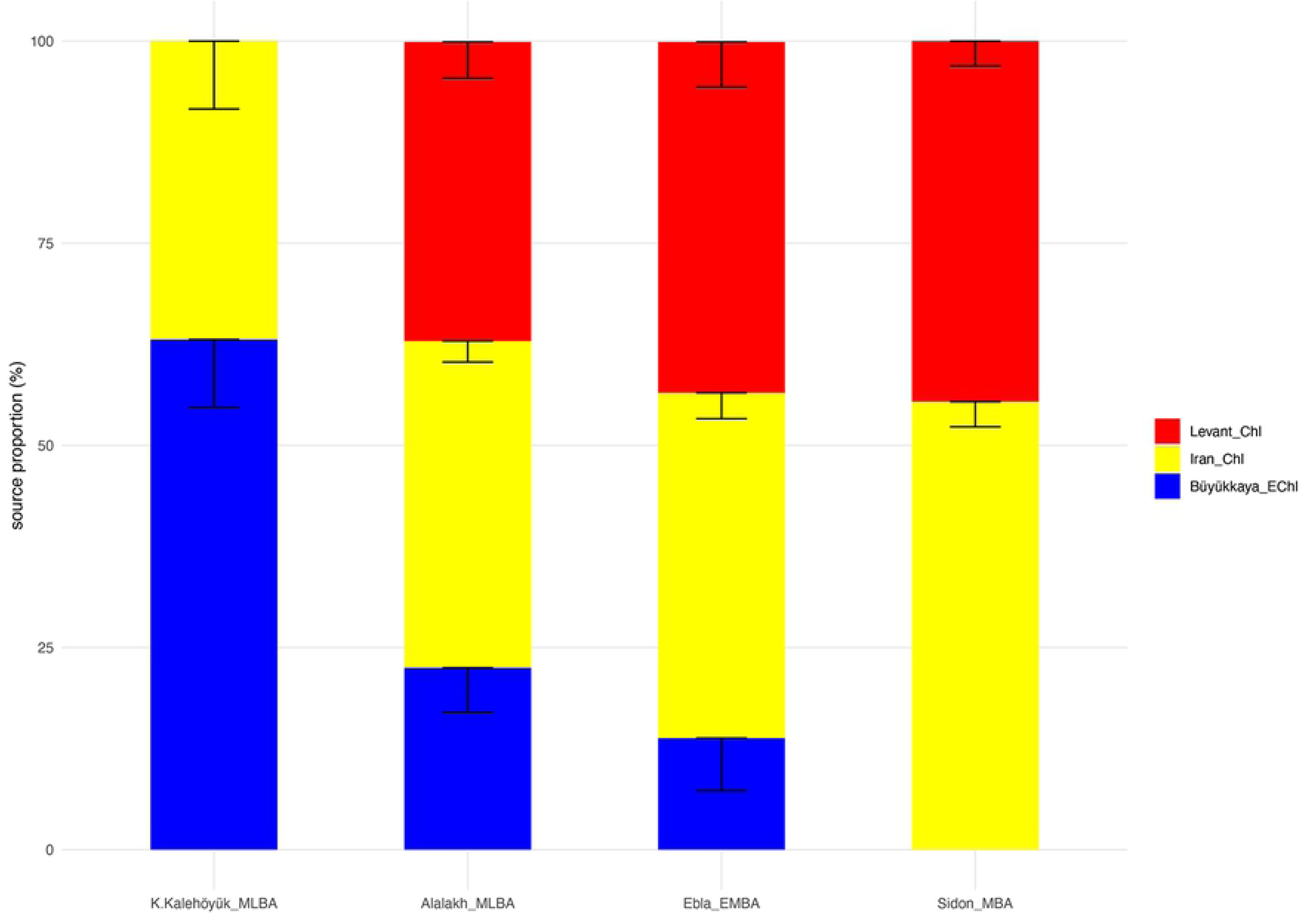
Admixture modeling (*qpAdm*) of Alalakh_MLBA, Ebla_EMBA, K.Kalehöyük_MLBA, and Sidon_MBA using Chalcolithic and Bronze Age source populations. Source proportions are plotted with -1SE. Abbreviations: E = early, M = middle, L = late, BA = Bronze Age, Chl = Chalcolithic.

Overall, these models provide adequate descriptions for the positioning of the individuals from Alalakh, excluding outlier ALA019, on the PCA in between contemporary Anatolian and central/southern Levantine individuals by breaking their ancestry down to three major components of Anatolian, Levantine, and eastern origin. For Alalakh, Ebla, and Sidon, models fit better with Iranian than Caucasus sources. However, when the Tell Kurdu population is used instead of Büyükkaya as a geographically proximal source, models with Caucasus sources fit better for Alalakh when Levant_EBA is used as a third source instead [46]. Therefore, a clear distinction between possible source populations from an eastern (Iranian) or northeastern (Caucasus) source is not yet possible with the data available. Sources to the east/southeast (northern and southern Mesopotamia) also need to be considered here, but these remain completely unsampled as of yet. The existing gaps in available genomic data touch on yet another important issue when performing admixture modeling: the individuals we group together here to represent ‘source populations’ need to be seen as mere proxies. We do not suggest that any of these groups are the actual source for admixture events. Indeed, based on archaeological and textual evidence, populations from northern Mesopotamia are among the likely genetic sources at Alalakh, especially the Hurrians and the Amorites, both groups known from texts to have been on the move in the region in the third and second millennia BC and which are attested in considerable numbers in the Alalakh texts [46, 53–55, 206–212].

#### Kinship analysis

READ computed on the total of 35 individuals from Alalakh successfully assigned pairs ALA011-ALA123 and ALA001-ALA038 as first degree related and pair ALA002-ALA038 as second degree related (Fig 14). The latter two cases are individuals from the Plastered Tomb and are reported in Skourtanioti et al. [46]. However, the genetic relatedness between ALA001 and ALA002 remains unresolved with this method, as the estimated P0 for this pair lies within the 95% confidence interval of the second-degree cutoff, but surpasses it in the +2 SE, and therefore either a second or higher degree are possible. Plotting *r* against k0 estimated by *lcMLkin* clusters pairs in three main groups that correlate with the result of READ: pairs ALA011-ALA123 and ALA001-ALA038, pairs ALA002-ALA038 and ALA001-ALA002, and all the other unrelated pairs (*r* ≈ 0) (Fig 15). For all related pairs, *r* is lower than expected, as suggested by the comparison with the degrees assigned by READ and by *r* = 0.9 between two different genomic libraries generated from the same individual (ALA039). Underestimation of *r* can be attributed to the lower quality of ancient data and has been reported before in Mittnik et al. [45], where genetic relatedness was explored in a large set of ancient individuals. However, the clustering of pairs ALA002-ALA038 (*r* = 0.16) and ALA001-ALA002 (*r* = 0.12) indicates that the latter most likely also represents a second-degree relationship. Interestingly, the two first-degree pairs ALA011-ALA123 and ALA001-ALA038 have both *r* = 0.39 but differ in the k0, and hence, suggesting a sibling-sibling and a parent-offspring relationship, respectively.

**Fig 14.**
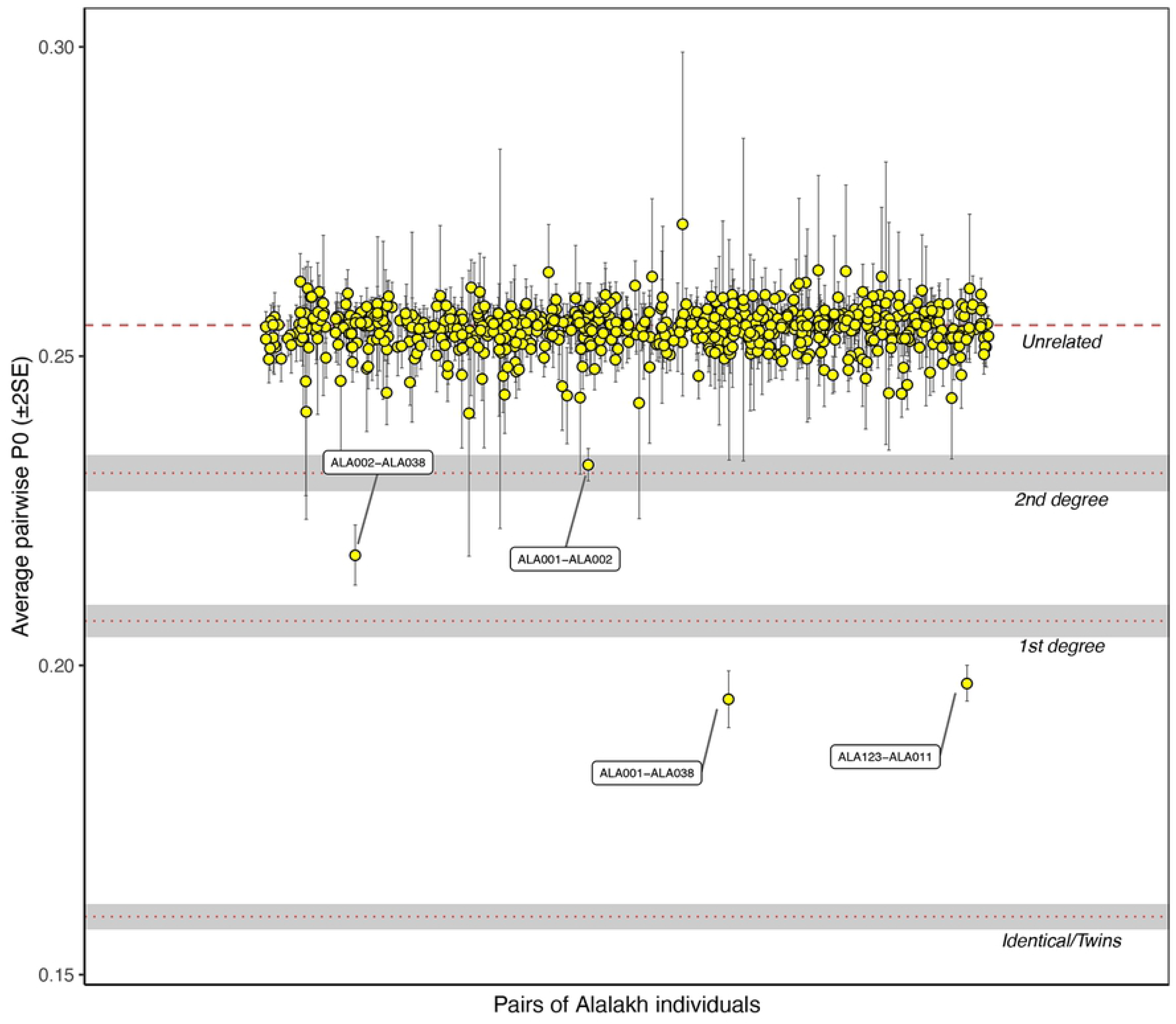
Kinship analysis with READ

**Fig 15.**
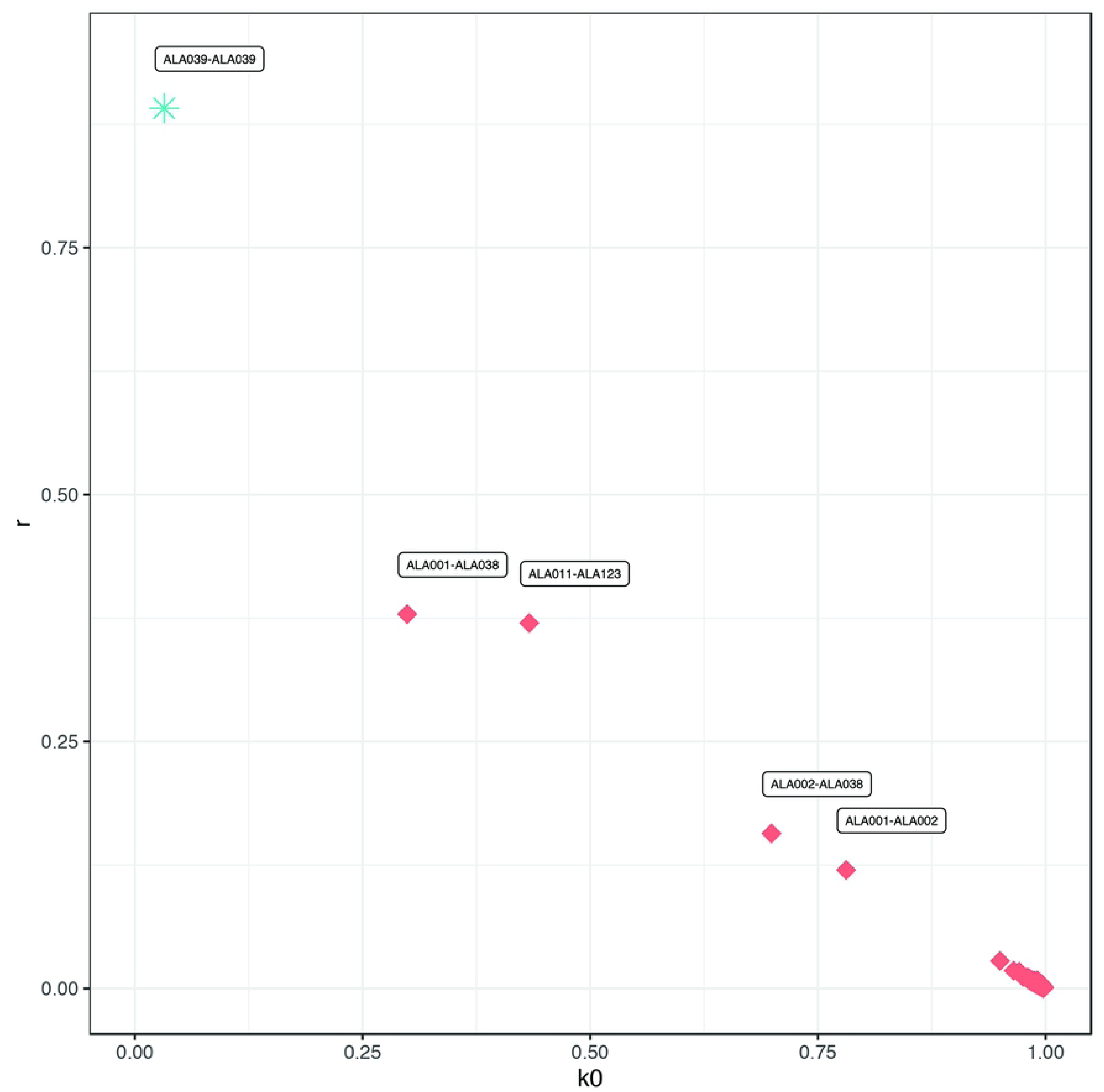
Kinship analysis with *lcMLkin*

Altogether, therefore, kinship in the first and second degree can be securely identified between five individuals from Alalakh. In all cases, the deceased were buried in close spatial proximity to one another. Individuals ALA011 and ALA123, two small children who were buried next to each other inside a casemate of the Area 3 fortification wall [76, 84] are first degree relatives, making them direct siblings. The other three individuals come from the Plastered Tomb and are discussed further below.

## Discussion

The aDNA analysis from Tell Atchana revealed that the sampled individuals are genetically very homogeneous – with the exception of ALA019 – and that the common ancestry at Alalakh was widespread over a larger area which stretched southeastward at least until Ebla. Consequently, aDNA’s resolution for scenarios of micro-regional migration might be limited. The genetic homogeneity of the samples from Alalakh suggests that the recent ancestors of most individuals came from within the wider Amuq-Ebla region, rather than beyond, which conforms well with the overall strontium and oxygen isotopic results that indicate a local upbringing within the Amuq Valley for the majority of sampled individuals.

Though the oxygen isotopic results are relatively homogenous, the strontium results are generally more informative. These suggest an overall population structure at the site during the MBA-LBA that was made up of a majority of people from the city itself. Based on the ancient faunal samples from Alalakh, we estimate that 40 individuals came from the city itself. The modern snail shells revealed that strontium ranges for many other locations within the Amuq Valley are comparable to those from Alalakh. This means we need to reckon with the possibility that a substantially larger portion of people than the eight that fall outside the Alalakh range but within the range calculated for the Amuq Valley originated within the Amuq Valley from sites other than Alalakh. Five individuals (9.4%) are identified as non-local to the whole Amuq Valley on the basis of the modern snail shells, one of which (ALA110) apparently moved to Alalakh during later childhood, resulting in the different ^87^Sr/^86^Sr ratios between M2 and M3 (see Fig 12, Table 4).

The only correlation between non-locals and any contextual variable such as burial location, type, or date, is the association of secondary burials with non-local individuals. One of these non-locals (ALA098) was found together as part of a secondary burial consisting only of three mandibles. The other two mandibles, ALA048 and ALA099, are as local to the Amuq, but not Alalakh. It therefore seems that all three of these individuals were born outside of Alalakh, although ALA048 and ALA099 seem to have grown up in the Amuq Valley. The wide separation between their ^87^Sr/^86^Sr values, however, indicates that all three spent their childhoods in different places (ALA098 = 0.706801; ALA048 = 0.707851; ALA099 = 0.707977), despite being buried together.

There are several potential explanations for this relationship between non-locals and secondary burials, not all of which are mutually exclusive. The most straightforward explanation is that these individuals moved to Alalakh at some point during their lives and then died and were buried there. If secondary burial was a stronger tradition in the area(s) where these individuals originally came from, it is possible that their families chose secondary burial for this reason, even though it was a minority practice at Alalakh itself [76, 77, 79]. However, given the nature of secondary burial, there are other possibilities. These individuals may have moved to Alalakh during their lifetimes, and, following their deaths, the majority of their remains could have been transferred back to their original settlement(s) for burial, with only parts of them remaining at Alalakh for burial. Alternatively, these individuals could have lived their entire lives elsewhere, but, after death, parts of the deceased could have been brought to Alalakh for burial, perhaps as a result of its status as a regional cult center [74, 75]. People who were able to do so may have chosen to inter a portion of their family’s remains at the cult center for a variety of reasons, including gaining favor from the gods, in order to raise their social standing, or because they were ritual specialists who were expected and entitled to do so.

Genomic data exists for only two (ALA037 and ALA004) of the five ^87^Sr/ ^86^Sr non-locals. Both individuals, ALA037 and ALA004, share the same genetic profile as the other individuals from Alalakh. There are two possible explanations for this pattern: both individuals could have come to Alalakh from a distance that is outside the Amuq but still within the wider Alalakh-Ebla catchment area, as the genomic data suggests, or this may be a case of backwards migration – the parents or grandparents of ALA037 and/or ALA004 could have emigrated from the area around the Amuq, ALA037 and ALA004 consequently spending their childhood elsewhere, but later coming back to Alalakh and subsequently dying there. As the ancestors of ALA004 and ALA0037 would have originated from the Amuq region in this scenario, their genetic profile matches the other individuals sampled from Alalakh.

### The case of the Well Lady (ALA019)

Aside from the bulk of genetic data from Alalakh that suggests regional ties over many generations, there is one outstanding case of long-distance mobility. Individual ALA019 – the Well Lady – takes up an extreme outlier position in the PCA closest to sampled individuals from Bronze Age Iran/Turkmenistan/Uzbekistan/Afghanistan, which can be confirmed with *outgroup f_3_* statistics [46]. While it is impossible to say exactly where to the east or northeast this individual (and/or her ancestors) came from, especially in the absence of data from nearby eastern regions like Mesopotamia, it is clear from the genetic data that either this individual or her recent ancestors migrated to the Alalakh region. The strontium isotope data allows us to narrow down the possibilities, and it seems that the Well Lady herself did not migrate, but rather her ancestors, as the ^87^Sr/ ^86^Sr ratios of all three molars sampled (M1, M2, and M3) fall within even the most narrowly defined local range for Alalakh (see Fig 12); however, due to a lack of research on bioavailable strontium isotopes in the Central Asian areas where the PCA suggests she came from, it is not currently possible to definitively rule out a childhood spent in these regions. A scenario in which she was part of a pastoral community that frequently came into contact with inhabitants of the Amuq Valley is unlikely, due to the low variation in all three ^87^Sr/ ^86^Sr values (M1 = 0.708456; M2 = 0.708474; M3 = 0.708540). The case of the Well Lady is therefore particularly interesting, not only because it is the only genetic outlier in a dataset of 37 individuals (if we add the Ebla data on top of that, in a dataset of 48 individuals), but also because the strontium evidence is consistent with her having spent her whole life at Alalakh; however, despite likely being a local of Alalakh, she did not receive a proper burial, instead found face down at the bottom of a well, with extremities splayed, indicating that she was thrown into the well.

The presence of this genetic outlier at Alalakh is generally not surprising, given the extensive genetic, archaeological, and textual evidence for long-distance contacts between both people and polities in the second millennium BC, and it is doubtful that she was the only such outlier present in the city throughout its history, especially considering that she herself was apparently not migratory. Indeed, dental morphology of the Well Lady shows shoveling of I2 [213], a feature which is passed down genetically and is shared by three other individuals – 42.10.130, buried in the Royal Precinct, ALA012, buried in the extramural cemetery, and ALA139, buried in the Area 4 cemetery – as well as ALA030 (the accident victim found in Area 3), ALA132, and ALA133 (both buried in the Area 4 cemetery), although the trait is less pronounced in these latter three individuals. Of these six individuals, only ALA030 has thus far yielded sufficient aDNA preservation, and this individual is not a genetic outlier among the Alalakh population. It is possible, therefore, that the former three individuals, which show pronounced I2 shoveling, may also be genetic outliers, similar to the Well Lady.

### The Plastered Tomb: evidence for local elites with kinship ties

The Plastered Tomb is the most elaborate, elite grave at Alalakh, judging from the grave construction and the richness of the burial goods [82]. While isotopic data could be generated for all four individuals in the tomb, genetic analysis only succeeded in three cases (ALA001, ALA002, and ALA038; ALA003 did not yield preserved aDNA), but this data illuminates the kinship ties between these individuals.

The four individuals buried in the Plastered Tomb were spatially arranged in three different layers atop each other, separated by plastering (Fig 16A). From a construction viewpoint, it is clear that the lowest two individuals, ALA001 and ALA003, were deposited first, and then the plastering over them was laid, sealing both bodies. On top, arranged above one another and separated by plastering, were put individuals ALA002 and ALA038. ALA038 was, furthermore, placed in a wooden coffin (unpreserved, but attested by wood impressions in the plaster surrounding it) [77, 82]. While this general order of interments is clear [82], the time interval between each burial is not – there could have been between one to up to four separate events; the semi-disarticulated state of ALA003’s remains [80] suggests that even the lowest two individuals may not have originally been placed in the grave at the same time.

**Fig 16.**
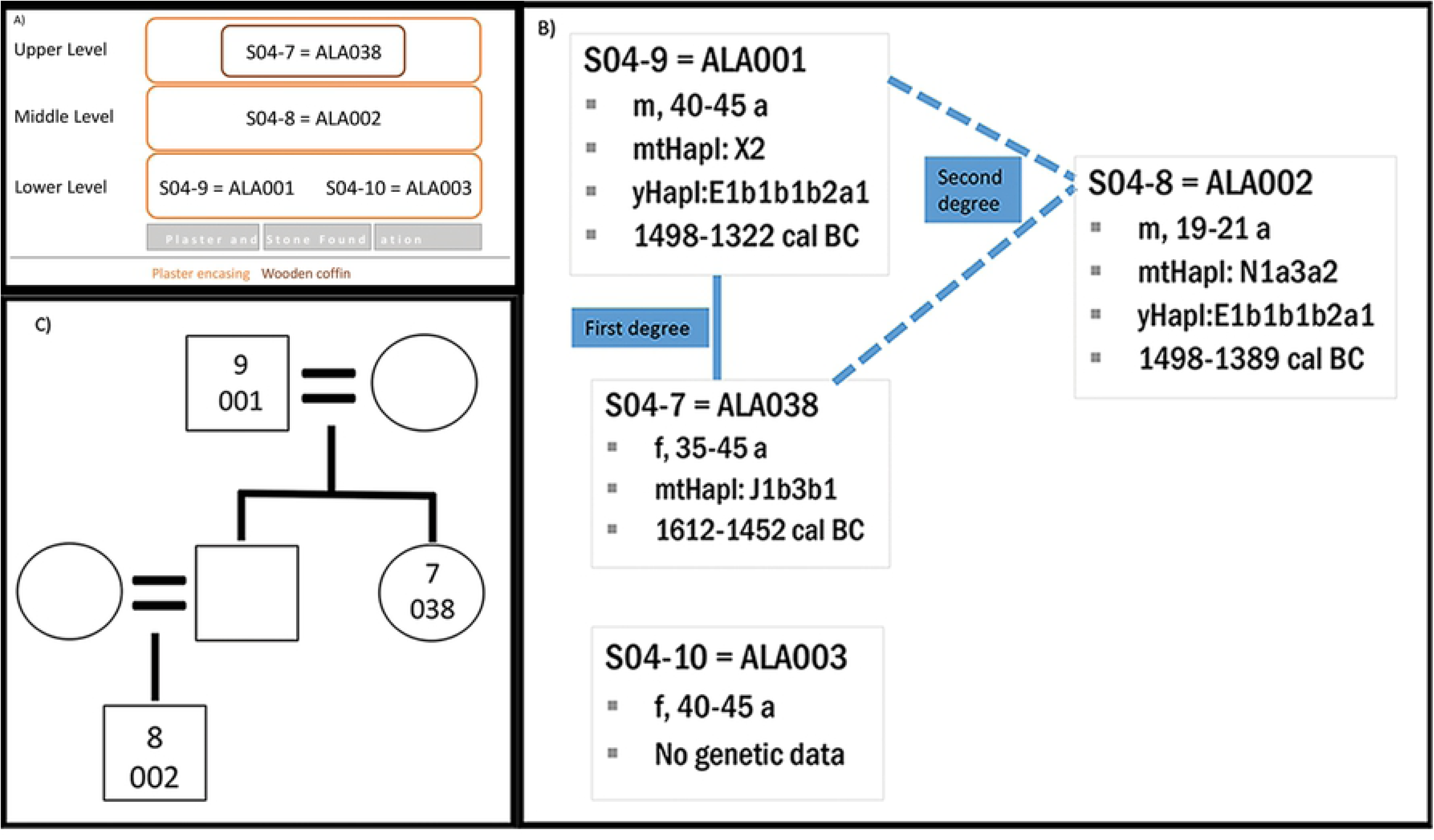
The Plastered Tomb: A) schematic representation of the spatial setting of the four individuals within the grave after Yener [82]; B) osteological and genetic information of the Plastered Tomb individuals, including biological kinship; C) family tree.

Osteological analyses concluded that three individuals in the grave were likely female and one individual (ALA001) male. ALA002 was tentatively ascribed as female on the basis of pelvic and cranial morphology and post-cranial robusticity [80, 81]. Genetic sexing has now revealed that this individual was actually male, which changes the arrangement of the tomb to an even sex ratio (2:2) [46]. According to the most recent analysis by R. Shafiq, the male individuals were estimated to have died at an age of 40-45 years (ALA001) and 19-21 years (ALA002); the two females were between 40-45 (ALA003) and 35-45 (ALA038) years old at death.

Multiple burials are common in the whole Levantine and Mesopotamian area during the MBA and LBA and are often associated with family burials, so even before genetic analysis, it was expected that these four individuals were related in some way [82]. The genetic data confirms, on the basis of READ [193] and *lcMLkin* [194], that all three successfully DNA sequenced individuals were biologically related (Fig 16B) [46]. None of them share the same mitochondrial haplogroup, which is exclusively passed on from mother to offspring. This means that first-degree relatives ALA001 and ALA038 are father and daughter, confirming the k0-based distinction of *lcMLkin* from siblings ALA011-ALA123. ALA002 must therefore be the nephew of ALA038 and the grandson of ALA001, linked to ALA001 via the male line, as they do not share the same mt-haplogroup but have the same Y-haplogroup (Fig 16C).

Stratigraphically, the tomb belongs to Period 4 at Alalakh and can be dated on the basis of the grave goods to the 15^th^ century BC [82, 214]. The radiocarbon dates of ALA001, ALA002, and ALA038 all confirm this dating. Furthermore, the combination with the kinship and osteological data enables a more precise dating: the overlap in date ranges from 1498-1452 BC between ALA001 and ALA038 – father and daughter, and both adults in their thirties or forties at their age of death – can be used to place them more precisely in time: both must have died during the first half of the 15^th^ century BC. The death of the grandson/nephew ALA002 would then be at the very latest during the first decades of the second half of the fifteenth century BC.

Examining these individuals as a group on a population genomics level shows that they cluster together with all other individuals from Alalakh and Ebla, excluding the Well Lady. Isotopic analysis confirms that ALA001, ALA002, and ALA038 likely grew up at Alalakh, while the difference in the strontium ratio of the two samples from ALA003 could indicate that this individual moved to Alalakh from within the Amuq Valley during early childhood (if an M1 was sampled by Meiggs [128]). Although it was not possible to generate genetic data for ALA003, her presence in the lower layer of the tomb and the semi-disarticulated state of her remains [80] suggest that she was also a part of this family group and was likely either from the same generation as ALA001 (perhaps his wife and/or his sister?) or an earlier one (perhaps his mother?). There are therefore selected members of at least three, possibly four, generations of a local, elite family buried in this unusual tomb that was so richly constructed and appointed and would have been so prominent outside the city wall [77, 82] – a unique tomb constructed for, and likely by, local elites as a potent symbol of their social status and power.

## Conclusions

Our investigation of the burial corpus at Alalakh via strontium and stable oxygen isotopic analysis, combined with both published [46] and new aDNA results, sheds light on aspects of human mobility at an urban center in the northern Levant during the MBA and LBA. The various lines of evidence reveal that most individuals grew up locally, with different levels of mobility, from long-distance to regional, indicated for a smaller number of individuals. We used overlap in datasets to refine signals for mobility, most notably by limiting the likely distance of the migrations. The strontium isotope data, due to its better refinement in outlier identification than the stable oxygen isotope data and to the different level it operates on than the aDNA data, proved to be best-suited for estimating numbers of non-locals and was even able to reveal that the Well Lady, though a remarkable genetic outlier, may have been local to Alalakh. Long-distance migration of the type demonstrated by this individual’s ancestors appears (at least from the data currently available) to be rather rare.

The arising picture from Alalakh’s population with regard to mobility is complex and cannot be easily paralleled with certain burial traditions, with the exception of secondary burial, which is associated with non-local individuals. As the case of the Well Lady indicates, though, we may be missing entire portions of the population due to their non-recovery for a variety of possible reasons. This example highlights how the vagaries of discovery and issues of representativeness influence mobility studies, and it is important to keep in mind that only a small portion of the total number of ancient inhabitants of the city has been recovered to date and is available for sampling. Nevertheless, this study has revealed multiple scales and levels of mobility at Alalakh in the Middle and Late Bronze Age, and shows, as have other recent studies in the ancient Near East [139, 141, 215], that the majority of sampled individuals were locals who likely lived, died, and were buried in close proximity to the place where they were born. This has important implications for understanding individual mobility in the Near Eastern Bronze Age: while such mobility is documented at relatively high levels both textually and archaeologically, it seems that – within the range and limitations the methods discussed here are able to determine – relatively few individuals were buried away from their homes. The majority of cases of long-distance mobility may therefore have been on a temporary basis, for the duration of a diplomatic mission or a specific crafting commission, for example, rather than permanent relocations.

## Acknowledgments

The analyses presented here were carried out under the auspices of the Max Planck-Harvard Research Center for the Archaeoscience of the Ancient Mediterranean (MHAAM) and the European Research Council’s (ERC) European Union’s Horizon 2020 research innovation programme (ERC-2015-StG 678901-Food-Transforms). Oxygen isotope analysis was conducted at the Max Planck Institute for the Science of Human History (MPI-SHH) in Jena, Germany under the supervision of P. Roberts and J. Ilgner; strontium isotope analysis was conducted at the University of Cape Town under the supervision of P. J. Le Roux; and aDNA analysis was conducted at MPI-SHH under the supervision of J. Krause, G. Brandt, and A. Wissgot. All contextual data, including site maps, is courtesy of the Alalakh Excavations project, and all skeletal data is courtesy of R. Shafiq. All necessary permits were obtained for the described study, which complied with all relevant regulations.

## Author contributions

Tara Ingman: Conceptualization, data curation, formal analysis, investigation, methodology, project administration, visualization, writing – original draft, writing – review & editing Stefanie Eisenmann: Conceptualization, data curation, formal analysis, investigation, methodology, project administration, visualization, writing – original draft, writing – review & editing

Eirini Skourtanioti: Methodology, Conceptualization, data curation, investigation, Writing – Original draft Preparation, Writing – Review and Editing

Murat Akar: Conceptualization, Writing – Review and Editing Jana Ilgner: Investigation, Data Curation, Validation

Guido Alberto Gnecchi Ruscone: Investigation, Data Curation Petrus le Roux: Investigation, Data Curation, Validation

Rula Shafiq: Data Curation, Investigation, Resources Gunnar U. Neumann: Investigation, Resources Marcel Keller: Investigation, Resources

Cäcilia Freund: Investigation, Data Curation, Validation Sara Marzo: Investigation, Data Curation, Validation Mary Lucas: Investigation, Data Curation, Validation Johannes Krause: Funding Acquisition, Supervision

Patrick Roberts: Conceptualization, Supervision, Investigation, Validation, Methodology, Writing – Review and Editing

K. Aslıhan Yener: Conceptualization, Supervision, Writing – Review and Editing

Philipp W. Stockhammer: Conceptualization, Funding Acquisition, Supervision, Writing – Review and Editing

## Text supplements

S1 Text. The chronology of the burials: detailed analysis of stratigraphy and radiocarbon dating.

S2 Text. Discussion of the modern snail shells and their underlying geology.

S3 Text. Comparison between modern environmental ^87^Sr/ ^86^Sr ratios in Meiggs [128] and the samples in this study.

## Table supplements

S1 Table. Isotopic results of all individuals, with sampled tooth indicated.

S2 Table. Admixture modeling results.

